# Assessing the impacts of agriculture and its trade on Philippine biodiversity

**DOI:** 10.1101/861815

**Authors:** Andrea Monica D. Ortiz, Justine Nicole V. Torres

## Abstract

The Philippines is home to a high number of unique species that can be found nowhere else in the world. However, its unique species and ecosystems are at high risk because of habitat loss and degradation. Agricultural land use and land use change are major drivers of biodiversity loss in the Philippines.

In the Philippines, an important area that requires focus is plantation agriculture (monocropping) for high-value crops such as banana and pineapple, which are grown widely in the country, particularly in the island of Mindanao. The intensive nature of plantation agriculture means that it has many adverse effects on the environment while producing goods and commodities that are typically for trade and export with international partners. This means that local biodiversity losses may be driven by countries thousands of kilometers away.

While many global studies have attempted to understand how biodiversity impacts are embodied within agricultural goods, there are few studies that have investigated the Philippines specifically. In this study, local and national-scale data are investigated to better characterize the nexus between agriculture, biodiversity, and trade in the Philippine context. Based on geographical data, many banana and pineapple plantations and their buffer zones interact and overlap with areas that are high in biodiversity, such as Protected Areas and Important Bird Areas. In this study, data shows that 82 threatened species, including the critically endangered Philippine eagle, are at risk of exposure to agricultural activities from high-value crops banana and pineapple. An additional and important political and legal analysis is also undertaken in the study to reveal key legislation and enabling environments relevant to the interactions between land use and biodiversity. More stringent definitions and protections for biodiversity are recommended to recognize the increasing role that agricultural production, and importantly, its global trade, has on threatened Philippine species and habitats.

## 1. Introduction and background

### 1.1 Philippine biodiversity, agriculture and land-use

The Philippines is a mega-biodiverse country, with nearly half of its approximately 1100 terrestrial vertebrates and vascular plants endemic to the archipelago (Posa et al., 2008). However, decades of poor natural resource management, overexploitation, and human activities have led to critical declines in biodiversity in the Philippines. High rates of deforestation during much of the century have decreased forest cover from about 70% in 1910 to less than 25% in 2003 (Kastner & Nonhebel, 2010; Kastner, 2009). More recent estimates put current Philippine forest cover at 3-5% (Posa et al., 2008).

Agricultural land use and land use change (LULUC, i.e. forest clearing to make land available for cultivation) were identified as the main drivers of biodiversity losses in a global study (Chaudhary et al., 2016). In particular, land use for rice, coconut, rubber, and palm oil production in Southeast Asian countries was found to contribute the most to biodiversity loss (Chaudhary et al., 2016).

Agricultural land is used to produce commodities for domestic consumption as well as for international trade and export. The global liberalization of trade means that products from the Philippines are exported to international partners thousands of kilometers away. The international trade of agricultural products can provide significant economic opportunities, as it enables the ability to consume ‘exotic’ or seasonal goods year-round, and benefits from the lower production costs in other countries (Fader et al., 2013). Many of these agricultural exports are high-value crops, such as bananas and pineapples, which are grown widely in the Philippines.

However, intensively grown crops like banana and pineapple have large adverse impacts on the environment, biodiversity, soil, and human health (e.g. Hernandez & Witter, 1996; Henriques et al., 1997; Hartemink, 2005; Billeter et al., 2007; Levillain et al., 2012; Echeverría-Sáenz et al., 2012; Shaver et al., 2015; Jadin et al., 2016; Chopin et al., 2016). Therefore, while trade facilitates the growing global demand for agricultural commodities -- typically from faraway, industrialized countries (e.g. Fader et al., 2013; Chaudhary & Kastner, 2016) -- products have ‘embodied’ negative biodiversity impacts.

For example, it has been found that agricultural exports from the Philippines to the United States rank eighth in terms of largest embodied biodiversity impacts globally (Chaudhary & Kastner, 2016). Bird extinction risks in the Philippines are also high due to tropical wood exports, and forests in Palawan island in particular are at high risk for biodiversity loss (Nishijima et al., 2016; Chaudhary & Mooers, 2018). These threats to biodiversity mean that the Philippines has significant challenges to meet increases in biomass demand without putting even higher strains on the ecosystems, particularly in light of population growth and dietary changes (Kastner & Nonhebel, 2010).

Given the importance of biodiversity conservation towards Sustainable Development Goals (e.g. SDG 15, Life on Land), as well as global conservation targets (e.g. the Aichi Targets of the Convention on Biological Diversity) vis-à-vis the increasing demand for food by a growing population, it is crucial to better understand biodiversity losses due to the interactions between agriculture and trade in the Philippines.

#### 1.1.1 Research focus

While a number of studies (e.g. Chaudhary & Kastner, 2016; Nishijima et al., 2016) have investigated the interlinkages between trade, agriculture and biodiversity with mention of the Philippines, they focus on the global level, and thus do not provide spatially explicit information on the species risks and implicated agricultural and forest goods that are linked to them. This limits the usability of these results to inform national or local conservation policy. In addition, it is also important to identify existing local legal frameworks that relate to this important nexus of agriculture, biodiversity, and trade to identify gaps and opportunities in protecting Philippine biodiversity.

A better characterization of species risks related to domestic agricultural production and its international trade is therefore needed to provide information on both biodiversity threats, as well as the interlinked policies that can enable better protection. In this paper, this need is addressed through an analysis of agriculture, trade, and biodiversity data. In addition, this assessment is contextualized within the political-legal landscape of the Philippines. Characterizing these threats and policies may contribute to the state of knowledge on the interactions between trade, agriculture and biodiversity, as well as provide evidence for strengthening or improving initiatives toward sustainable development amidst biodiversity losses and increased agricultural LULUC.

The analysis begins, firstly, with a review of the impacts of intensive, export-oriented agriculture, followed by a glimpse at the policies surrounding habitat protection in the Philippines.

### 1.2 Impacts of intensive agriculture on the environment

A review of intensively grown cropping systems reveals that monoculture production has a myriad of negative impacts on the environment. Banana and pineapple are typically cultivated in monocrop (single-crop) plantations, and in the Philippines, mostly in provinces on the island of Mindanao. There are a number of definitions of plantation agriculture, but it is often thought of as a large-scale, mostly foreign-owned, export-oriented, high-input/high-output farming system (Hartemink, 2005). For example, bananas are usually produced in large plantations with fixed infrastructures and high inputs of fertilizers, pesticides, and irrigation (Ploetz et al., 2015).

Because of their design, plantations increase the concentration of food sources for other organisms, including pests and diseases, which then multiply quickly and can affect the harvest (Hernandez & Witter, 1996). High-production systems already require high levels of input (e.g. fertilizers) to produce the commodity, but then also need high levels of control inputs (e.g. pesticides) to eliminate competing organisms (Hernandez & Witter, 1996). These high quantities of inputs can result in chemical leaching, surface runoff, erosion, poor soil fertility, and emissions that have significant negative impacts on the environment through contamination of terrestrial and aquatic ecosystems (Hernandez & Witter, 1996; Henriques et al., 1997; Hartemink, 2005).

Pesticide residues from banana and pineapple plantations have been found to lead to fish killings and water quality problems (Echeverría-Sáenz et al., 2012). They have also been documented to cause serious health consequences for both humans and wildlife, particularly birds (Hernandez & Witter, 1996; Billeter et al., 2007). Runoff and manufacturing processes (e.g. washing of bananas prior to shipping) also provide entry for toxic substances into water (Henriques et al., 1997). Pests and soil organisms may also gain resistance to pesticides and other chemicals, leading to more use of inputs (Hernandez & Witter, 1996).

In terms of biodiversity, plantations contain a low diversity of wildlife as compared with natural forests. Plantations restrict natural habitat and favor only a very restricted number of co-habiting species (Hartemink, 2005). They also simplify and homogenize the landscape, which has negative effects on biodiversity by reducing forest cover, increasing the isolation of native species in remnant forest, impeding the movement of species across landscapes, and increasing habitat fragmentation (Shaver et al., 2015). Furthermore, plantations of pineapple often do not respect safety vegetation margins, which makes agricultural impacts on ecosystems even greater (Echeverría-Sáenz, 2012).

This means that agricultural intensification (e.g. through plantations) is a major driver of global biodiversity loss (Tscharntke et al., 2005). However, it has been argued that a larger threat than these adverse impacts is the loss of habitat through agricultural expansion (Hartemink, 2005). In the future, the combined effects of both agricultural intensification and expansion will have serious consequences for biodiversity (e.g. Chaudhary & Mooers, 2018; Kehoe et al., 2017; Delzeit et al., 2017). One of the ways to mitigate habitat loss and its negative effects is forest conservation. In the next section, the policies on Philippine Protected Areas are reviewed.

### 1.3 Philippine Policy on Protected Areas

Republic Act 7586, or the National Integrated Protected Areas System (NIPAS) Act of 1992, governs the establishment and management of Protected Areas in the Philippines. This policy was recently updated by Republic Act 11038, or the Expanded NIPAS (E-NIPAS) Act of 2018. Under these laws, the national Protected Areas system “encompasses the ecologically rich and unique areas and biologically important public lands that are habitats of rare and threatened species of plants and animals, biogeographic zones and related ecosystems, whether terrestrial, wetland or marine (E-NIPAS Act 2018 Section 2).” The Protected Areas system may be made up of areas classified as National Parks, Natural Biotic Areas, Natural Parks, Natural Monuments, Protected Landscapes and Seascapes, Critical Habitats and Wildlife Sanctuaries. Each category reflects the area’s size, natural conditions, physical features and purpose.

At the time of its passage, the NIPAS Law was considered a “regional leader in collaborative management of Protected Areas (Parr, 2017),” owing to its provision for multi-stakeholder participation through the Protected Area Management Board (PAMB) and Protected Area Management Plan (PAMP). The PAMB exercises broad oversight, management and monitoring powers. In addition to its mandate to review and approve policies, plans and activities within, and pertinent to the Protected Area, it is likewise authorized to allocate resources to implement the PAMP, manage the Protected Area incomes and set user fees and charges (E-NIPAS Act 2018 Section 11a). It is a multi-stakeholder body, with representation from national government agencies, local government units, local civil society organizations and indigenous peoples communities (NIPAS Act 1992 Section 11). The E-NIPAS Act has since expanded this membership to include sitting Senators who reside in the Protected Area’s locality, as well as representatives from local academic institutions and the private sector (E-NIPAS Act 2018 Section 11).

The PAMP serves as a ten-year framework for the Protected Area’s administration. The E-NIPAS Act provides guidance on what this strategy should contain, including, among others, zoning, buffer zone management, habitat conservation and rehabilitation, research and policy development (E-NIPAS Act 2018 Section 9). The preparation of the PAMP provides further opportunities for multi-stakeholder engagement, as participation from Indigenous Peoples and local communities, Civil Society and the private sector is specifically provided for, notably in the determination of the zoning regime, development of interventions for each zone and enforcement (DENR, 2019).

#### 1.3.1 Protected Areas in the Philippines

At present, the Philippines has recognized 244 Protected Areas (PAs). In total, these add up to 7.76 million hectares of terrestrial, wetland and marine ecosystems (DENR-BMB, 2018). Ninety-four of these PAs were recently recognized under the E-NIPAS Act, but had been earlier identified and delineated by various Executive actions.

In 2014, the Philippine Biodiversity Management Bureau and the German Agency for International Cooperation (GIZ) conducted an assessment of 64 Philippine PAs. This evaluation found numerous issues confronting the areas studied, including “low buy-in” of nearby local governments and communities, and conflicts in land tenure, boundaries and zoning (Guiang & Braganza, 2014). PAMB operations were also found to be severely hampered by inadequate budgetary resources and personnel (Guiang & Braganza, 2014). As a result of these, and other challenges, the report noted that the PAs surveyed were largely unable to address threats to biodiversity and safeguard critical habitats and ecosystems (Guiang & Braganza, 2014).

The E-NIPAS Act and its Implementing Rules have made efforts to address the issues surfaced in the assessment. Apart from specifying clearer powers and functions for the PAMB, this amendment also provides for harmonization of the PAMP with other local sectoral plans (E-NIPAS Act 2018 Section 9). Typically, these include the local governments’ Comprehensive Land Use Plans and Ancestral Domain Sustainable Development and Protection Plans for PAs that may overlap with Indigenous Peoples’ territories. Because of the potential overlaps and interactions between agricultural LULUC and PAs, these policies that protect biodiversity and habitats are important to the research focus and objectives of the study.

### 1.4 Objectives of the study

In this paper, it is investigated how high-value agricultural crops, particularly pineapple and banana, have impacts on Philippine biodiversity through the analysis of agricultural, economic, and biodiversity data related to production and trade. It is hypothesized that these products, which are highly valued in international markets or are produced on a large-scale, are linked to a high number of species under threat.

The paper also outlines pertinent laws, policies and agreements that consider the interactions in the agriculture-biodiversity-trade nexus. This analysis is used to outline of potential gaps and opportunities in the field, and as basis for recommendations about how the complex feedbacks between agricultural production, species loss and international demand can be better considered in the Philippine legal context.

## 2. Data and methods

### 2.1 Agricultural data

Banana and pineapple were chosen for the focus of the study based on their high production, export value, and the large amount of land area dedicated to their cultivation. Data on their cultivation was obtained from the Philippine Statistics Authority (PSA) which has records until 2018. Banana varieties were not differentiated in production values, although Cavendish is the primary export variety. Agricultural data on planted area was used to determine which Philippine provinces were main producers (large-scale, to consider export volume) of the high-value crops. A threshold of more than 10,000 hectares was set to be considered a large provincial producer.

Additional trade data on production from the Department of Trade and Industry (DTI), PSA and agricultural company profiles were used to determine the foreign country destinations of banana and pineapple exports to link import demand and local production. Site plantation locations for banana and pineapple were obtained from the Bureau of Plant Industry (BPI), the agency responsible for accreditation of prospective growers of fruit and vegetable crops, and providing the corresponding documentation for them.

Geographical data used in the study included Philippine country and provincial boundary shapefiles obtained from the Database of Global Administrative Areas (GADM). OpenStreetMap (OSM) was used to obtain approximate latitude-longitude coordinates of these reported plantations at the barangay level. Plantation sites were not included where data was ambiguous (e.g. recorded barangay name did not match geographical records) or when available only at the more coarse-scaled municipality level.

To demonstrate the proximity of known plantations to environmentally sensitive areas, PA shapefiles were obtained from the Philippine GIS Data Clearinghouse and the World Database on PAs. A 10-kilometer buffer diameter was added around each plantation point to account for farm size and leakage/spillover effects from intensive agricultural production. This size of buffer was chosen to conform to the European Commission Digital Observatory for Protected Areas for the Agricultural Pressure Indicator (EC-DOPA, 2019).

### 2.2 Protected areas and buffer zones

These boundaries are considered in the analysis, as Philippine PAs rely on zoning as a strategy for management and administration. This zoning system determines the types of activities that are permissible within the PA. Strict protection zones are “closed to human activities by virtue of their significant biodiversity value, or high susceptibility to geo-hazards and other dangers (DENR, 2019).” Multiple use zones, on the other hand, refer to areas wherein “settlement, traditional and sustainable land use, including agriculture, agroforestry, extraction and activities and income generating or livelihood activities may be allowed to the extent prescribed in the PAMP (DENR, 2019).”

Delineation of the PA’s boundaries may also involve the determination of and regulation in immediately adjacent Buffer Zones, to provide the PA with additional safeguards against pollution, invasive alien species and human encroachment (DENR, 2019). These Buffer Zones are established along with the PA, and are identified based on ecological, economic and social criteria, as assessed by the PAMB and approved by the DENR Secretary (E-NIPAS Act 2018 Section 8, DENR, 2019). Once established, these areas are managed by the PAMB in coordination with other stakeholders and government units (DENR, 2019). Owners of private lands that may be encompassed by the Buffer Zones are likewise directed to develop their properties in accordance with the PAMP (DENR, 2019).

Historically, the formal establishment of Buffer Zones has not been without contention. In at least two documented cases, areas with geothermal energy developments were designated as protected area buffer zones, although these were well away from the peripheries and within what would otherwise have been considered the core “strict protection” zones (La Vina et al., 2009).

Apart from the identified ecological, economic and social criteria, current guidelines do not specify further prescriptions for the assessment and identification Buffer Zone. As a result, the extent and measurements of established Buffer Zones varies widely among PAs.

### 2.3 Biodiversity data

Species occurrence data were obtained from the Global Biodiversity Information Facility (GBIF) which contains hundreds of millions of records obtained from observations, scientific records and specimens, to citizen science data collections (GBIF, 2019). Species records from the Philippines were selected to obtain a species occurrence list for land fauna (mammals, birds, reptiles and amphibians). Species data was cross-referenced to the threatened fauna species list from the Biodiversity Management Bureau of the Philippine Department of Environment and Natural Resources (DENR-BMB, 2017) to geo-locate threatened species occurrences.

A separate database containing geotagged vulnerable tree species (Ramos et al., 2012) was also included to account for threatened plant biodiversity. Marine biodiversity was not considered in the study because of the focus on land-based agricultural impacts, although agricultural LULUC has significant impacts on marine biodiversity from surface runoff (See Section 1.2).

To identify threatened species within agricultural plantations, firstly, key provinces for large scale banana and pineapple production were identified based on BPI and PSA data to find a geographical focus. PA boundaries, Important Bird Areas (BirdLife, 2018), plantation sites (with their 10-km buffer), and biodiversity data were overlaid using R statistical software to analyze interactions between where production is high and intensive, alongside where threatened species are distributed, and where areas have some level of protection. Species which had records of occurrence from GBIF within the buffer zone of plantations were identified. By doing so, these species are recognized as being affected by their proximity and range overlap with intensive agricultural production, a known threat to biodiversity.

## 3. Results

### 3.1 High value crop production

Based on PSA data, planted area of pineapple plantations in 2018 in two provinces—Bukidnon and South Cotabato—made up over 72% of all pineapple planted area in the Philippines: 48 thousand hectares (ha) of a total 66 thousand ha were dedicated solely to the crop. For banana plantations, Davao del Norte, Compostela Valley and Bukidnon held the majority of planted area, with more than 20 thousand ha per province. A number of other provinces also had more than 10,000 ha of planted area for banana. Apart from Iloilo, Isabela, and Southern Leyte, all provinces that were large banana producers are on the island of Mindanao. Because the scope of interest is international trade, Tables 1a-1b thus show the hectarage of planted areas for pineapple and banana in Mindanao, respectively.

**Table 1a.**
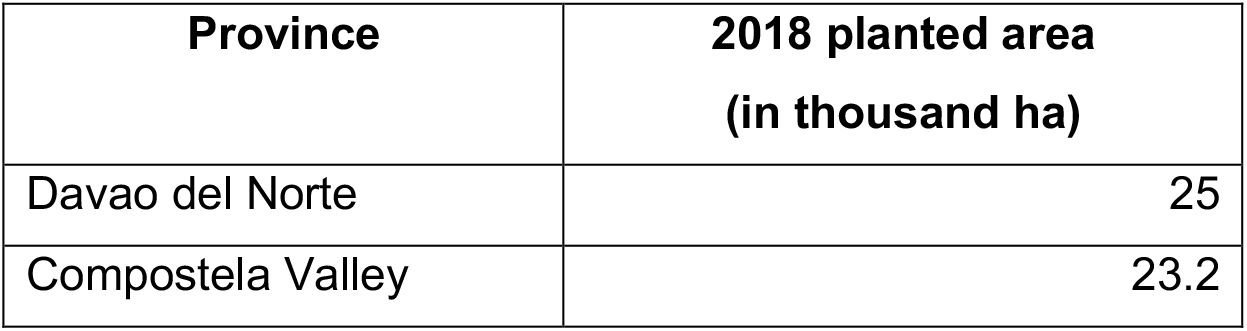
Large provincial producers: planted area, pineapple (Mindanao).

**Table 1b.** Large provincial producers: planted area, banana (Mindanao).

| **Province** | **2018 planted area**<br>**(in thousand ha)** |
| --- | --- |
| Davao del Norte | 37.3 |
| Compostela Valley | 21.8 |
| Bukidnon | 21.2 |
| Maguindanao | 18.2 |
| Misamis Oriental | 17.4 |
| Davao del Sur | 15.4 |
| Cotabato | 14.3 |
| Agusan del Sur | 11.2 |
| Davao Oriental | 10.8 |

### 3.2 Threatened species occurrence

#### 3.2.1 Country-level distribution of threatened species occurrence

GBIF data on species occurrence shows that many sites in the Philippines are host to high biodiversity, including various threatened species of fauna and trees. Many of the records of species occurrence can be found both within and outside of PAs (Figure 1). There are some observable positive biases in reporting, for example in the National Capital Region (NCR), and sparse data in some regions (e.g. Cordillera region, where there is an IBA) but these limitations are likely due to the nature of the collection of GBIF records, which is a well-documented limitation of the dataset (e.g. Beck et al., 2014).

#### 3.2.2 Focus on Mindanao agricultural plantations

Results of the analysis are centered around Mindanao because of its high level of agricultural land use and activity (see Tables 1a and 1b). Available BPI data was able to provide approximate barangay locations for 227 plantations for banana and 46 for pineapple (226 and 26 at barangay level, respectively). These were associated to 48 companies/groups for banana and 12 for pineapple. Although this may not be a complete figure of the number and scale of plantations in Mindanao -- particularly for pineapple which had more limited data compared to banana -- the data showed a good match between the high-production provinces and the geolocation of these individually identified plantations (See Figure 2 and Appendix).

Many of the plantations dedicated to pineapple and banana are oriented toward production for export by large multinational companies. Analysis of the DTI trade data showed that for bananas, most of the harvest is sent to Chinese markets, while the largest volume of pineapples is currently exported to Japan (Tables 2a and 2b).

#### 3.2.2 Interactions with biodiversity

A map of approximately geolocated banana and pineapple plantations at the barangay level in Mindanao showed that there are many interactions between plantations, PAs, and Important Bird Areas (IBAs). For example, it can be observed that there are a large number of both banana and pineapple plantations in Bukidnon that overlap with species occurrence records, the Mount Kitanglad and Kalatungan PAs, and several IBAs. There were also a large number of overlapping boundary zones between plantations in the provinces of Davao del Norte, Compostela Valley and Davao del Sur, the majority of which are banana plantations. These plantations have direct interactions and potential overlaps between threatened species occurrence with the Aliwagwag Protected Landscape, Mainit Hot Springs Natural Park, and Mount Apo Natural Park (See Appendix for more information).

There were also numerous species-land use interactions in Cotabato, South Cotabato, Sultan Kudarat and Maguindanao with an overlap of the agricultural buffer zone with the Allah Valley watershed/natural areas. This means that there are acute threats to biodiversity, including birds in IBAs, from the loss of natural habitat from plantation-style (monocrop) agriculture. With more complete plantation data, examples of these interactions between agricultural LULUC and biodiversity are likely to be observed elsewhere in the country throughout the land use interface between plantations and natural habitat.

**Table 2a.** Major export destinations of Philippine pineapple.

| **Plantation** | **Pineapple Variety** | **Destination** |
| --- | --- | --- |
| Dole | Tropical Gold | New Zealand, Malaysia, Philippines, other Asia Markets |
|  | Sweetio (with crown) | Japan, Korea |
|  | Sweetio (no crown) | China |

**Table 2b.**
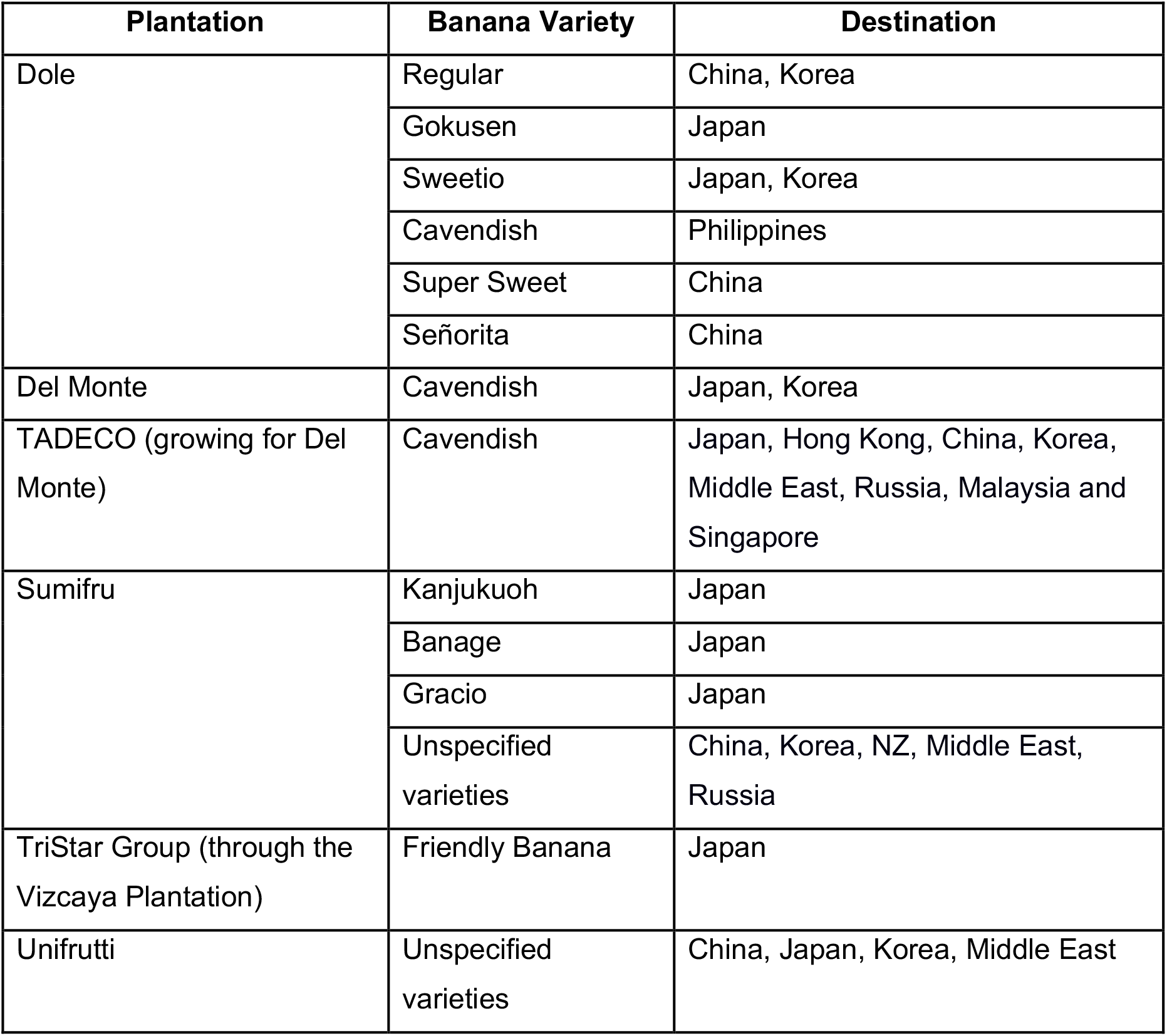
Major export destinations of Philippine bananas.

| **Plantation** | **Banana Variety** | **Destination** |
| --- | --- | --- |
| Dole | Regular | China, Korea |
|  | Gokusen | Japan |
|  | Sweetio | Japan, Korea |
|  | Cavendish | Philippines |
|  | Super Sweet | China |
|  | Señorita | China |
| Del Monte | Cavendish | Japan, Korea |
| TADECO (growing for Del Monte) | Cavendish | Japan, Hong Kong, China, Korea, Middle East, Russia, Malaysia and Singapore |
| Sumifru | Kanjukuoh | Japan |
|  | Banage | Japan |
|  | Gracio | Japan |
|  | Unspecified varieties | China, Korea, NZ, Middle East, Russia |
| TriStar Group (through the Vizcaya Plantation) | Friendly Banana | Japan |
| Unifrutti | Unspecified varieties | China, Japan, Korea, Middle East |

**Figure 1.**
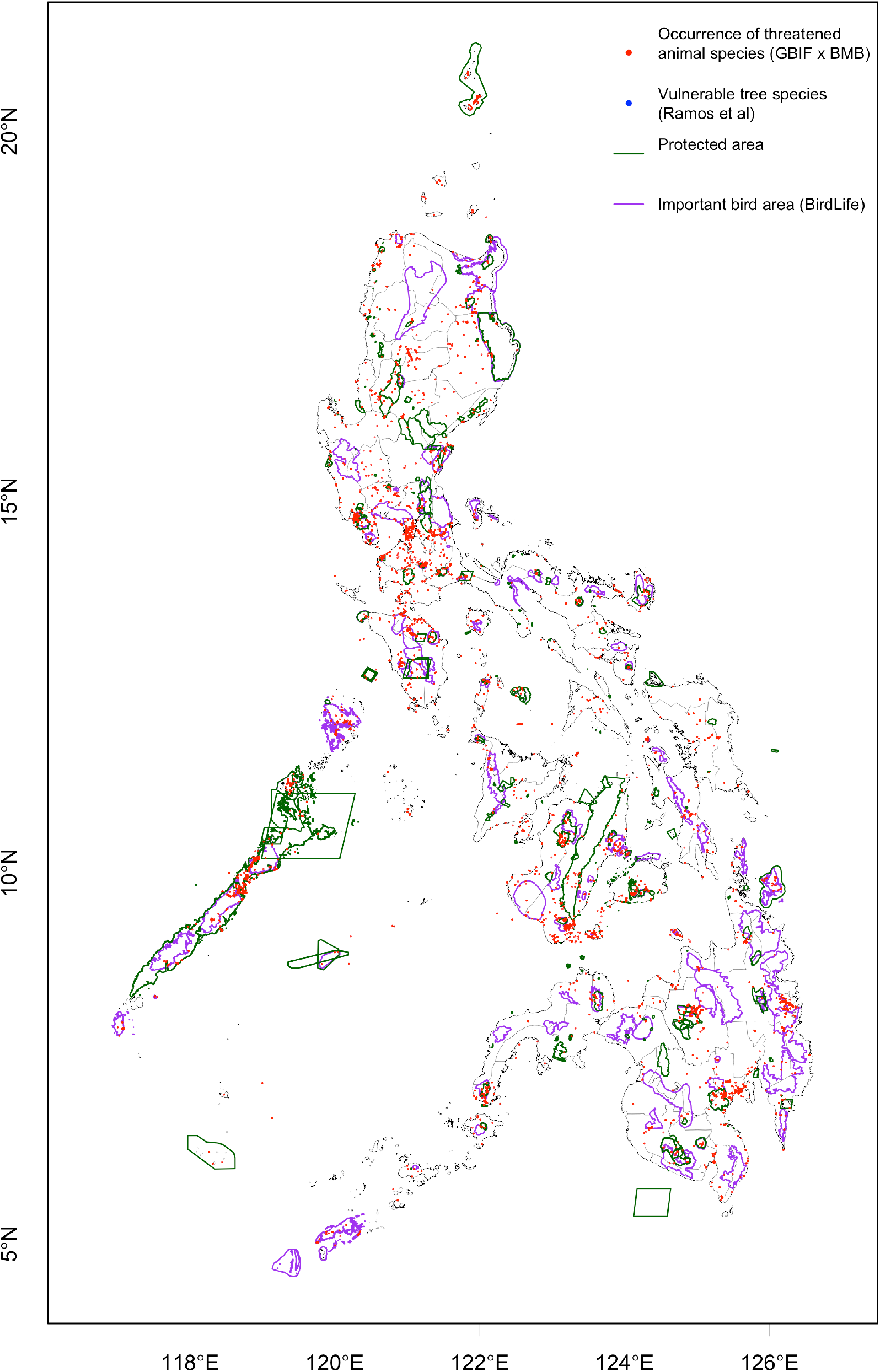
Occurrence of DENR-BMB listed threatened fauna from GBIF database and vulnerable tree species (from Ramos et al., 2012). Protected areas are outlined in dark green and Important Bird Areas in purple.

**Figure 2:**
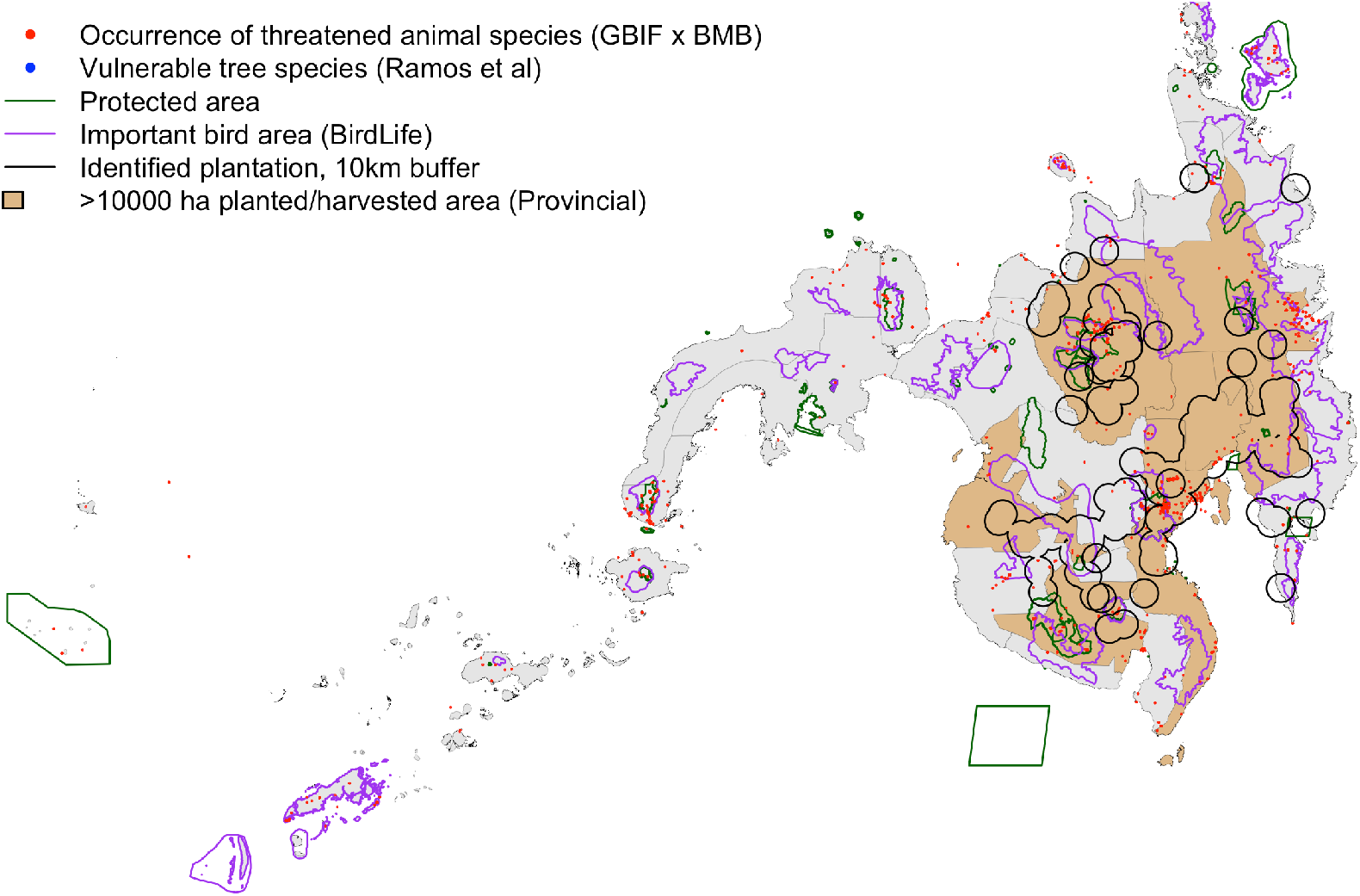
Large provincial producers of high value crops of banana and pineapple in Mindanao. Points indicate threatened species occurrence (tree species and fauna), and circles show identified pineapple and banana plantations with a 10-km buffer.

#### 3.2.3 Species threats linked to plantations

Based on the buffer zones and species occurrence records surrounding pineapple and banana plantations (Figure 2), there are 82 direct species threats for fauna (55 species) and vulnerable tree species (27) (Table 3a). Due to the numerous overlaps with IBAs (See Fig. 2), it is unsurprising that of the 55 threatened animal species, 47 of them are birds, which are important indicators of biodiversity (Blair, 1999; Pereira & Cooper, 2006; Schulze et al., 2004). Some of these are critically endangered species according to the DENR-BMB (2017), including the Philippine eagle and Peregrine falcon. Among mammals, there are records of the critically endangered Philippine naked-backed fruit bat occurring in these agricultural areas.

A large number of endangered birds are also noted to have occurred within agricultural land-use buffer zones, particularly from the *Accipitridae* family, which encompasses large birds including eagles, harriers, hawks, and kites. These have been noted in GBIF to have occurred within the buffer zones. Some species from the *Psittacidae* (true parrot) family are also endangered, and have occurred in these agricultural areas and their buffer zones. Several endangered owl species are also noted to have occurred in the buffer zones.

A number of threatened tree species from Ramos et al. (2012) are also found within the agricultural buffer zones of the noted pineapple and banana plantations, including the critically endangered *kamagong (Diospyros blancoi)*, and endangered tree species such as the *molave (Vitex parviflora)*, and *tindalo (Afzelia rhomboidea)*, among others (Table 3b).

**Table 3a.**
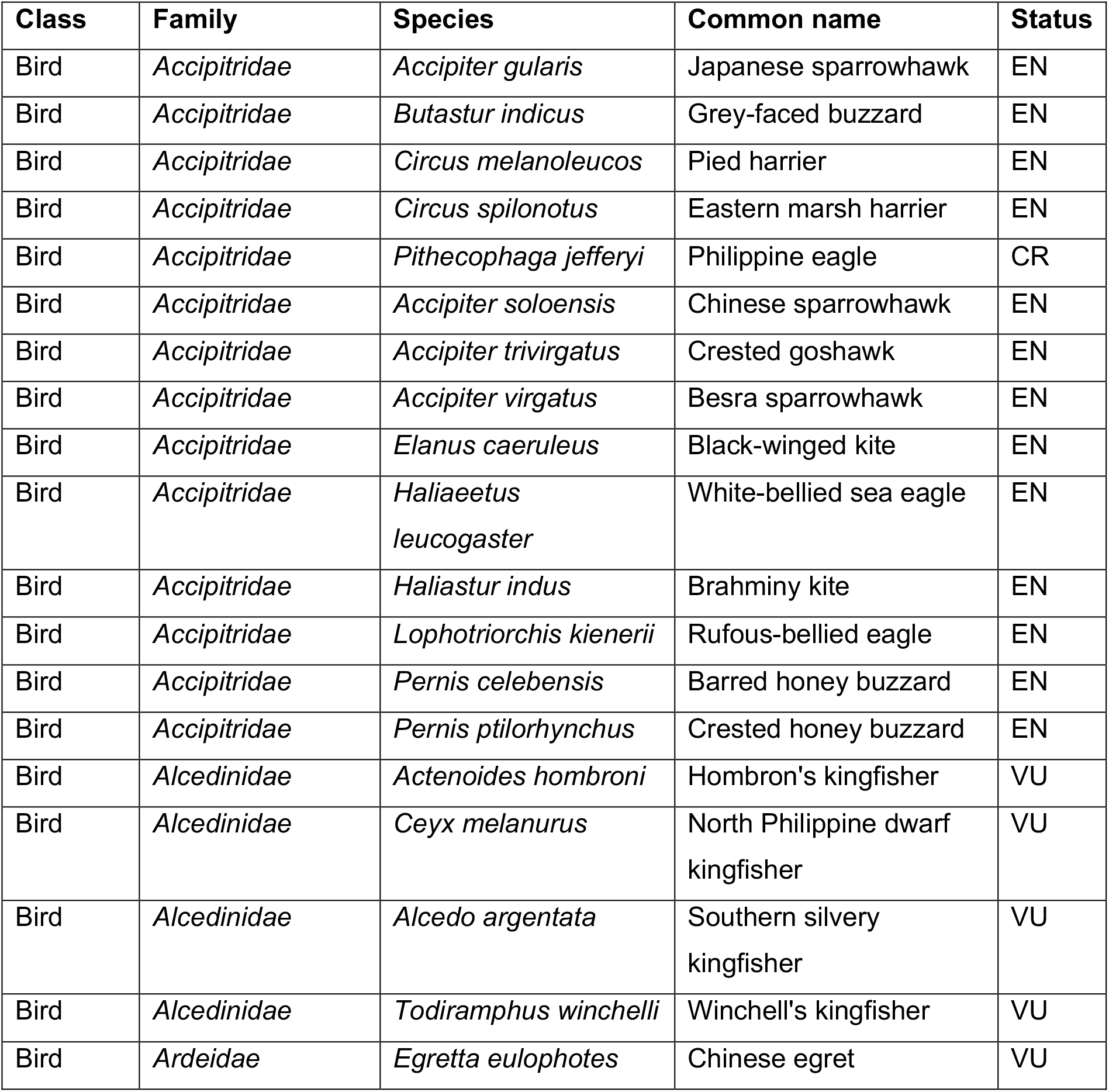
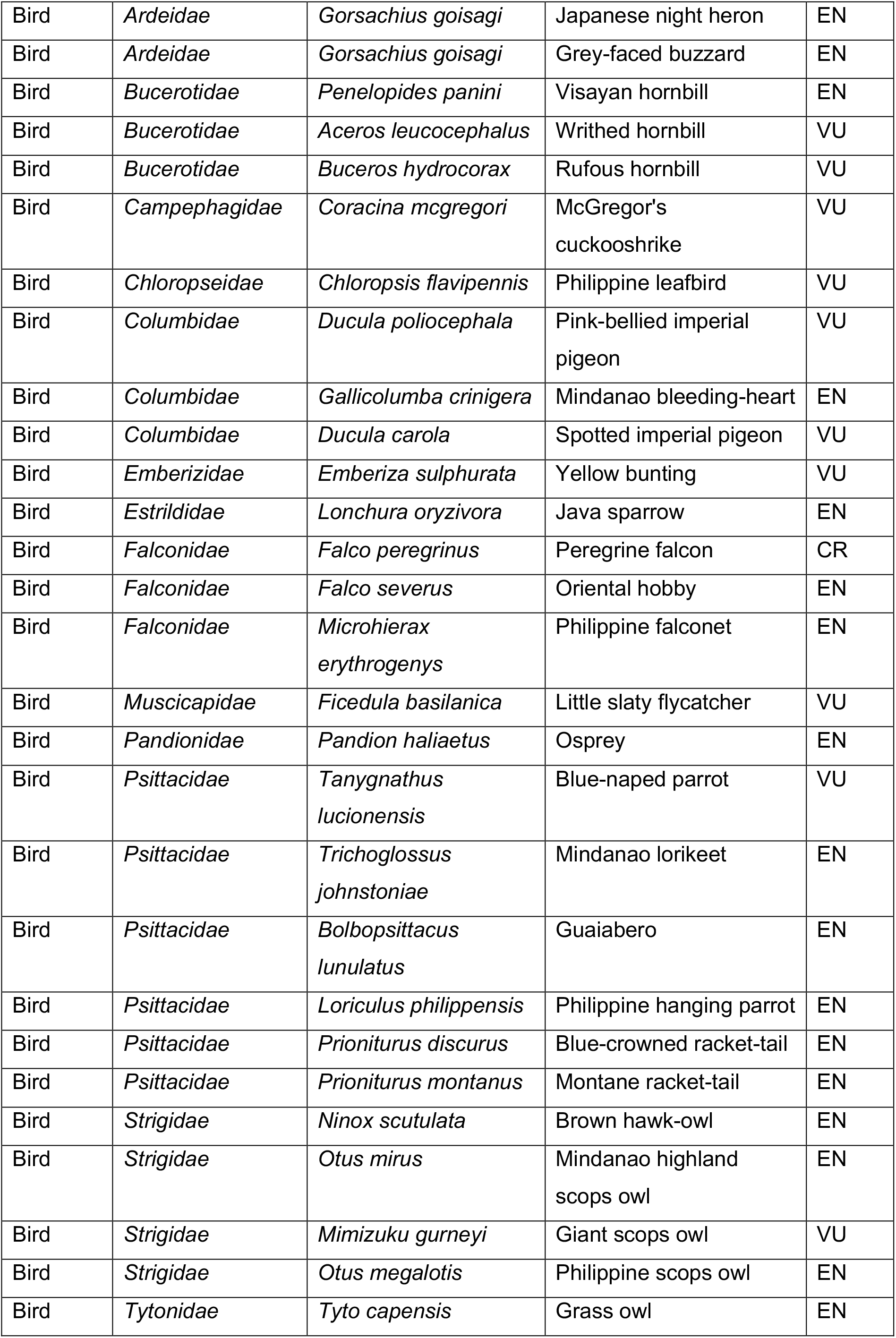
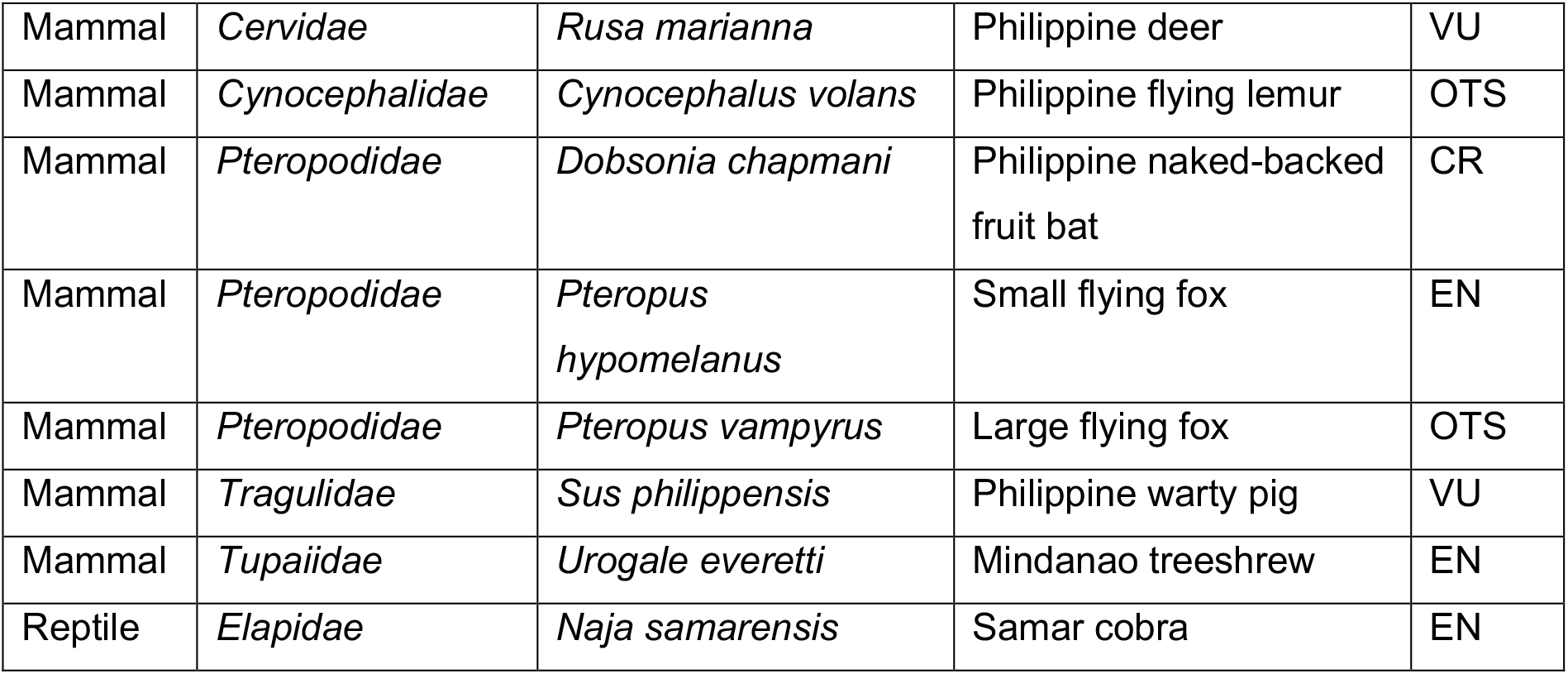
Threatened animal species within agricultural area and its buffer zone. CR is critically endangered, EN is endangered, VU is vulnerable, and OTS is other threatened species.

**Table 3b.** Threatened tree species (from Ramos et al., 2012) within agricultural area and its buffer zone. CR is critically endangered, EN is endangered, VU is vulnerable, and OTS is other threatened species. n/a indicates no information from the source study.

| Family | Species | Status |
| --- | --- | --- |
| *Anacardiaceae* | *Koordersiodendron pinnatum* | VU |
| *Araucariaceae* | *Agathis philippinensis* | VU |
| *Burseraceae* | *Canarium ovatum* | OTS |
| *Cannabaceae* | *Celtis luzonica* | n/a |
| *Dipterocarpaceae* | *Dipterocarpus gracilis* | VU |
| *Dipterocarpaceae* | *Dipterocarpus grandiflorus* | n/a |
| *Dipterocarpaceae* | *Dipterocarpus validus* | n/a |
| *Dipterocarpaceae* | *Hopea acuminata* | CR |
| *Dipterocarpaceae* | *Hopea plagata* | EN |
| *Dipterocarpaceae* | *Parashorea malaanonan* | n/a |
| *Dipterocarpaceae* | *Shorea almon* | VU |
| *Dipterocarpaceae* | *Shorea astylosa* | CR |
| *Dipterocarpaceae* | *Shorea guiso* | n/a |
| *Dipterocarpaceae* | *Shorea palosapis* | n/a |
| *Dipterocarpaceae* | *Vatica mangachapoi* | VU |
| *Ebenaceae* | *Diospyros blancoi* | CR |
| *Ebenaceae* | *Diospyros curanii* | VU |
| *Ebenaceae* | *Diospyros pilosanthera* | EN |
| *Fabaceae* | *Adenanthera intermedia* | OTS |
| *Fabaceae* | *Afzelia rhomboidea* | EN |
| *Fabaceae* | *Sindora supa* | EN |
| *Lauraceae* | *Cinnamomum mercadoi* | VU |
| *Meliaceae* | *Aglaia rimosa* | VU |
| *Meliaceae* | *Aphanamixis polystachya* | VU |
| *Meliaceae* | *Toona calantas* | CR |
| *Sapotaceae* | *Palaquium luzoniense* | VU |
| *Verbenaceae* | *Vitex parviflora* | EN |

### 3.3 Political and legal context of agricultural trade and biodiversity

An analysis of pertinent national policies and plans reveals that there are several instruments and issuances that relate to the interactions between agricultural production, its trade, and the environment. Firstly, while agriculture is an important industry in the country, the Philippine Development Plan for 2017 to 2022 (PDP) acknowledges the poor performance of the agricultural sector, even as it accounts for a large share of employment in rural areas (NEDA, 2017). To address this in the coming years, the government seeks to expand opportunities in the agricultural industry by improving productivity and enhancing commercialization (NEDA, 2017).

The PDP points out the economic potential of high-value crops such as bananas and pineapples the rapidly expanding export market, noting that production of these markedly increased from 2013 to 2015 (NEDA, 2017). Responding to this, interventions were proposed around the goal of diversifying production of selected agricultural commodities in specific regions of the country. Bananas, in particular, were identified as a potential high-value crop for Northern Mindanao (NEDA, 2017).

The PDP also lays out strategies for the Environment and Natural Resources sector. However, although these emphasize the need for a ridge to reef approach and sustainable integrated area development (NEDA, 2017), it is unclear how these might factor into the agricultural targets. This is especially significant, because while crop diversification and intensified commercialization are touted to deliver significant economic benefits, these may simultaneously involve trade-offs in terms of conservation goals.

Under the Philippine Environmental Impact Statement (EIS) System, agricultural plantations are not automatically considered Environmentally Critical Projects that must comply with the guidelines on Environmental Impact Assessments (EIA) and secure an Environmental Compliance Certificate (ECC) before operations may commence. Nevertheless, plantations may become subject to the EIS system if they fall within the enumerated Environmentally Critical Areas, such as declared national parks, watershed reserves and wildlife sanctuaries, habitats for threatened species of indigenous flora and fauna, and classified prime agricultural lands, among others.

The Revised Procedural Manual for the EIS System provides further clarity on the application of these regulations. Under these instructions, it is the size of an agricultural plantation in an Environmentally Critical Area that determines the type of process for prospective proponents to comply with. Plantations spanning 1,000 hectares or more must comply with the full EIS process, while smaller areas may undergo an abbreviated review with less comprehensive documentation (DENR-EMB, 2007).

EIAs and ECCs would be requirements at the outset of a plantation development. For those already in operation, the Bureau of Plant Industry (BPI) in 2017 outlined a Philippine Code for Good Agricultural Practice (PhilGAP) for Fruit and Vegetables Farming, which draws from the GAP standards adopted by the ASEAN Member States in 2006. Compliant farms and growers are issued corresponding Certification by the BPI. This confirms that they “address environmental, economic and social sustainability which result in safe and quality agricultural products (BPI, 2018),” entitling them to use the official PhilGAP markings for a two-year period. Many large plantations for banana and pineapple crops have been certified under this policy.

## 4. Discussion

### 4.1 Impacts on threatened species

In this study, the analysis of biodiversity, agriculture, and trade data has shown that the demand for agricultural commodities such as banana and pineapple has significant impacts on Philippine biodiversity. Firstly, the review of impacts of monoculture plantations demonstrates that the intensive nature of these fruit plantations concentrates chemical loads and utilizes a large amount of resources which can have direct and indirect impacts on biodiversity due to contamination and habitat loss (See Section 1.2). The land dedicated to these plantations has significant impacts on biodiversity due to the direct impacts associated with agricultural intensification, but also because the growing global demand for products could result in significant agricultural expansion and habitat loss.

Secondly, the geographical analysis shows that although many individually identified plantations may not occur directly within PAs, many plantations’ buffer zones overlapped or were in close proximity to recorded locations of many threatened species and to PAs and IBAs. Within buffer zones, some of these impacts (e.g. surface runoff of pesticides, effects of boundaries and habitat fragmentation) may be carried over to harm wildlife and habitats. Some threatened species themselves occurred within the boundary zones of the plantation sites. Many of these species are endangered endemic birds, which are known to be sensitive to chemical inputs used in plantations (Hernandez & Witter, 1996; Billeter et al., 2007), making the presence and proximity of plantations within sensitive and known IBAs a direct threat to biodiversity, particularly to threatened Philippine bird species.

A limitation of the study is that the number of identified plantations from BPI, while detailed in spatial scale, may not necessarily be reflective of the true number of banana plantations, including contract growers or smallholder farms (e.g. de la Cruz & Jansen, 2018) which may supply larger banana companies on smaller, fragmented pieces of land. Despite this, these identified plantations and their buffer zones are examples of how different land use types interact with records of species occurrence – a proxy for species presence and distribution – can already be observed in Mindanao. Although it is not analyzed in this study, systemic monitoring of threatened species over time could reveal whether there are significant correlations, and even causation, between plantation expansion and declines in species richness and abundance based on their occurrence.

### 4.2 Biodiversity connections with agricultural trade

In this paper, data showed that most species threats connected to pineapple and banana products are driven by international trade partners China, Japan, and South Korea (Figures 3a-b). For banana, China is the main export partner (over 1.1 million tons in 2018) while Japan is the main recipient of pineapple grown in the Philippines (over 150,000 tons in 2018).

It has been observed that China as the primary market for Philippine bananas is a recent development, attributed by analysts to the current administration’s efforts at strengthened political and economic ties with Beijing (Venzon, 2019). Aside from this political landscape, export tariffs and sanitary and phytosanitary conditions may account for this shift. Under the Philippines-Japan Economic Partnership Agreement (PJEPA), a bilateral treaty that entered into force in 2008, Philippine bananas are currently levied tariffs of anywhere from 2.5 percent to 18 percent, depending on the season (JPEPA Annex I).

Japan is also stringent as regards compliance with sanitary and phytosanitary conditions; for example, in 2018, Japanese authorities required testing for all bananas from the Philippines, after the presence of chemical insecticide Fipronil was found to be in excess of food safety standards (Banzon, 2019). In contrast, under the Tariff Reduction Schedule of the ASEAN-China Free Trade Agreement (ACFTA), Philippine bananas exported to China have been subject to zero tariffs since 2012. Growers have also noted that the Chinese market is less restrictive in terms of sanitary and phytosanitary standards, as well as less particular about the fruits’ appearance and packaging (Venzon, 2019).

Despite the high tariffs and reduced exports, however, the Japanese market’s preference for higher-grade bananas reflects significantly in the higher prices that the fruit sells for. Data from the DTI in 2018 shows that although Chinese exports edged out Japan’s with a difference in quantity of almost 200,000 tons, this only translated to a little over USD 10 million in terms of value. Early figures from 2019 show a similar trend - between China and Japan, the value difference is less than USD 1 million, although China received almost 130,000 tons more in terms of quantity.

It is likewise interesting how the shift to the Chinese market has had a less conspicuous effect on pineapple exports. While under the ACFTA, pineapple exports from the Philippines also enjoy zero tariffs, trade duties for pineapples under the PJEPA are also considerably less stiff. Instead of a seasonal tariff rate, such as that imposed on bananas, pineapple exports are instead entitled to a tariff rate quota of 1,800 tons, provided that each fruit weighs less than 900 grams.

**Figure 3a:**
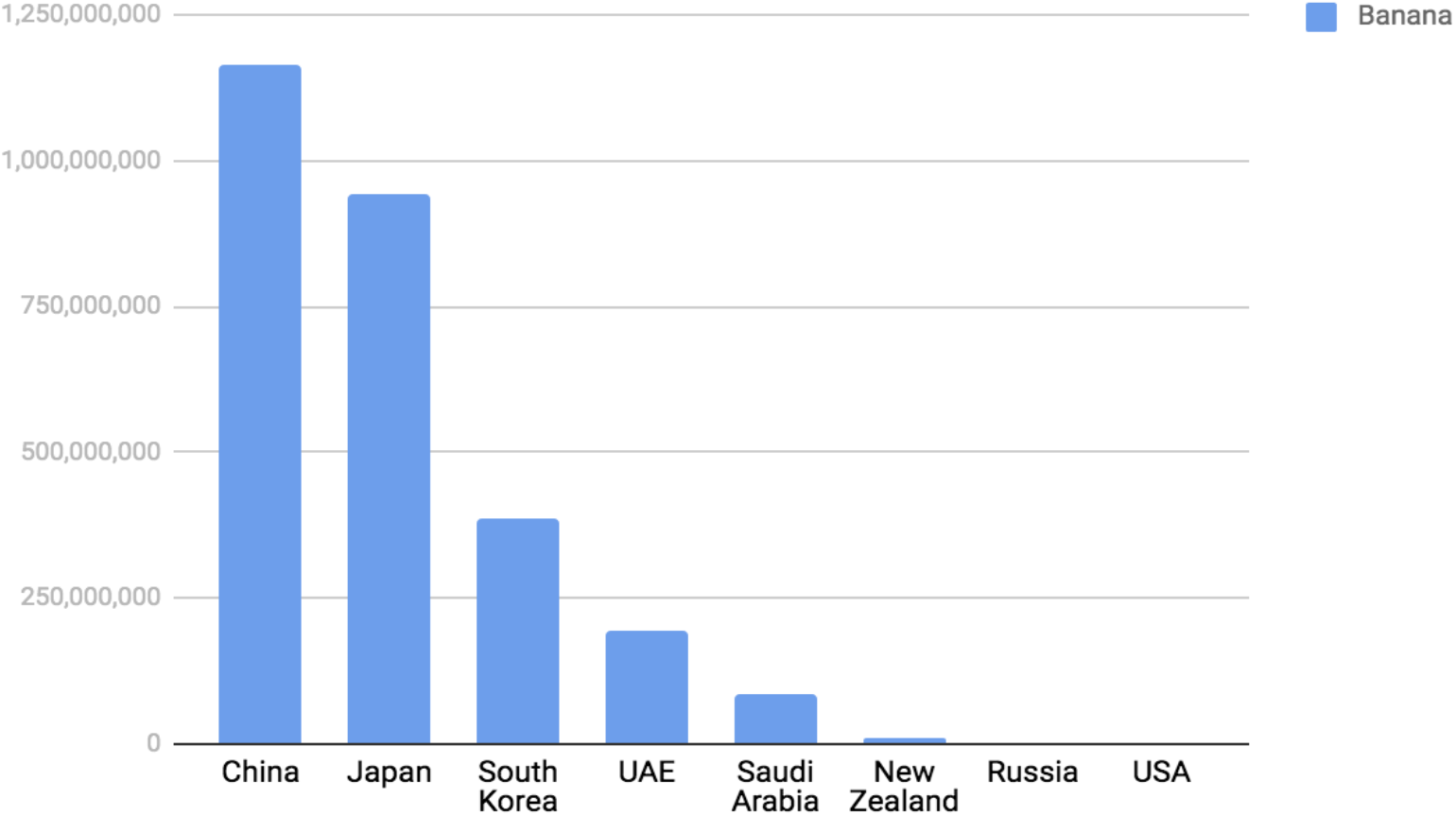
Quantity (in kg) of banana varieties/ products traded internationally, by country (DTI, 2018).

**Figure 3b:**
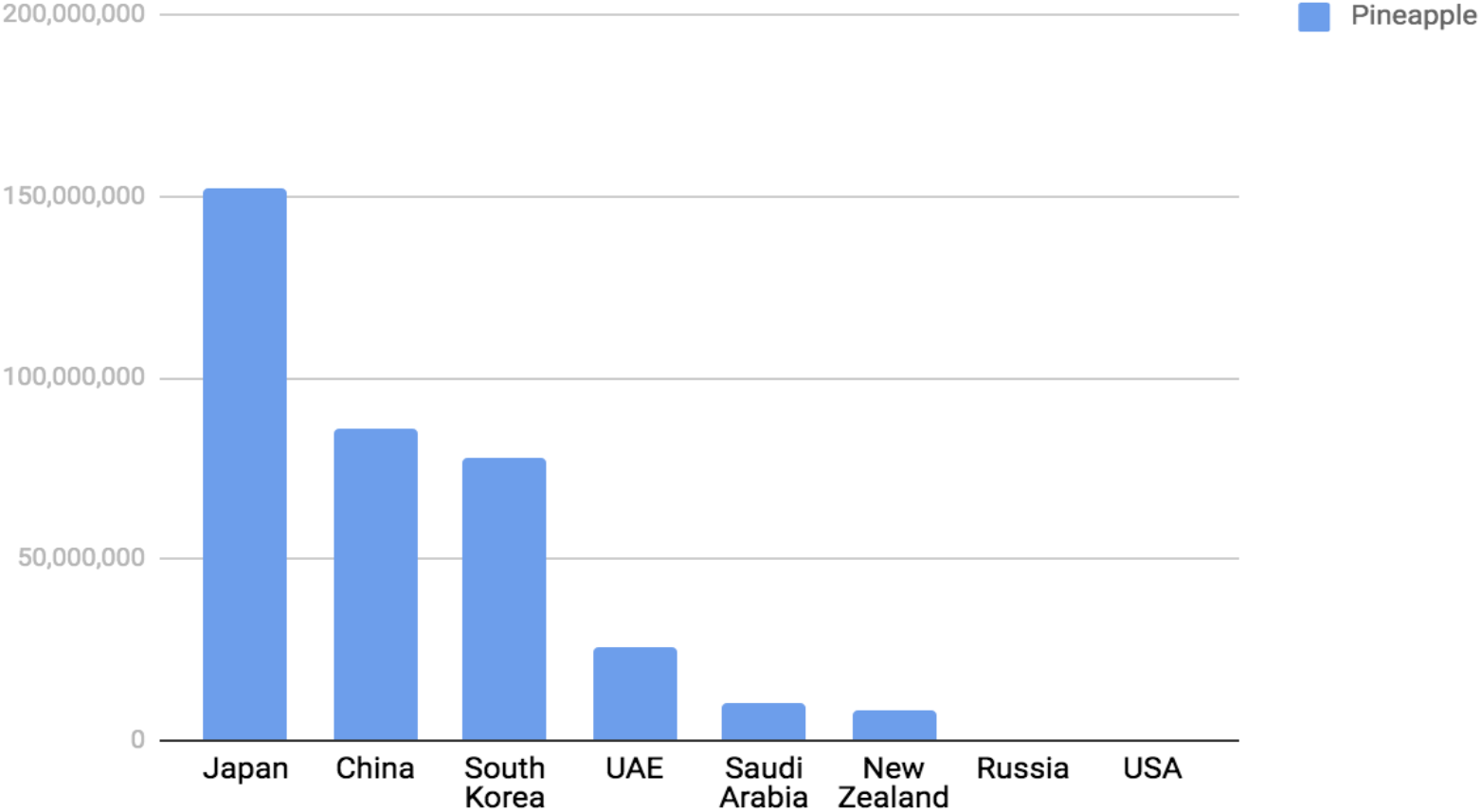
Quantity (in kg) of pineapple varieties/ products traded internationally, by country (DTI, 2018).

#### 4.2.1 The increasing role of trade

Reflecting upon these different export partners and the context of the political and economic exchanges between them and the Philippines emphasizes the role of trade in connecting agricultural supply, demand, and biodiversity impacts. Although the importance of trade in the context of biodiversity and agriculture is only recently becoming more understood, recent discourse has focused on how the spatial decoupling of food production and demand is increasing in recent years due to the ease and ability of trade to make connections between countries (Fader et al., 2013; Chaudhary & Kastner, 2016; Lenzen et al., 2012; Dalin & Rodríguez-Iturbe, 2016), which is only anticipated to grow in the future. Although the international trade of products may lead to benefits to the local economy, there are often trade-offs between economy and biodiversity, including increasing environmental pressures (Fader et al., 2013).

This means that international trade drives biodiversity threats, leaving large environmental and social footprints largely in developing nations that grow certain crops for export (Wiedmann & Lenzen, 2018; Lenzen et al., 2012). A well-known example of this is the China-Brazil soy trade, where lowering of tariffs that made trade easier led to increased deforestation of forests in Brazil to grow soy cheaply for export, a “teleconnection” that has led to significant impacts on biodiversity (e.g. Sun et al, 2017; Liu, 2014; Carrasco et al., 2017; Gibbs et al., 2015).

Therefore, the connection between who is demanding a product and where it is produced is significant -- because although the species threats may be local (e.g. the Philippine threatened endemic species reported here) -- the driving demand for intensively produced agricultural products is global, as demonstrated by the data on trade partners and the specialty exported varieties of crops (see Tables 1a-1b). Here, knowing the enabling policy environment and context (e.g. ASEAN tariffs, or Japan’s fruit quality standards in this study’s context) becomes important.

So while local plantations and agricultural intensification are responsible for the negative impacts that can affect native species, there is a concurrent and arguably greater need to look at the bigger picture of how biodiversity is affected by global trade. Biodiversity loss should be examined as a global systemic phenomenon, instead of looking at the degrading or polluting producers in isolation (Lenzen et al., 2012), particularly because biodiversity impacts are hidden in internationally traded food items (Chaudhary & Kastner, 2016).

Apart from looking at the connections between land use and consumption (e.g. Yu et al., 2013), it thus becomes important to inform consumers of the embodied impacts of products (Chaudhary & Kastner, 2016), a recent approach adopted by consumer-oriented groups dealing with the environmental impacts of palm oil through an eco-labelling effort (Ostfeld et al., 2019). However, there should be greater accountability not only on the demand side but also on the producers’ side. In this regard, there are several opportunities and conflicts with existing policy, or its lack thereof, in the context of the intersections between biodiversity and the agricultural trade of high-value crops in the Philippines.

### 4.3 Opportunities and conflicts with policy

Based on the reviewed key policies and legislation (See Sections 1.3 and 3.3), there are several points that are worthy of further discussion and investigation. Firstly, aims to expand and improve national productivity in agricultural production (e.g. in the PDP), particularly for high-value crops such as banana and pineapple, may come into conflict with biodiversity targets to stop biodiversity losses and protect habitats and ecosystems. As discussed in Section 1.2, agricultural expansion and intensification have detrimental impacts on biodiversity (e.g. Kehoe et al., 2017).

Therefore, recommendations to expand or intensify production to meet economic goals or targets need to be mediated by evidence-based discussion of their impacts on already dwindling natural habitat and forest cover in the Philippines. Mitigation of the damaging and long-lasting effects of intensive agriculture should also be considered vis-à-vis agricultural expansion/intensification and the international commitments of the Philippines to preserve and protect remaining habitats and biodiversity.

In this regard, there are opportunities to strengthen the existing framing of biodiversity in policies and legislation. The biodiversity safeguards currently provided for in the PhilGAP Code are very broad. Biodiverse areas are considered “sensitive areas”, necessitating additional management measures as regards waste water disposal and/or ground or aerial application of chemicals (BPI 2018). The Code however does not specify what form these measures should take, or what minimum prescriptions they should comply with.

The BPI had also earlier developed a Code for Good Agricultural Practice (GAP) for banana production specifically. These guidelines contain almost no biodiversity safeguards, save for a broad directive on the use of integrated pest management strategies to avoid possible ecological imbalance (BPI, 2013). Furthermore, it focuses in large part on more internal concerns, such as “microbiological, chemical and physical food safety risks during production, harvesting, and post-harvest handling and distribution (BPI, 2013).”

The national standards for pineapple production, on the other hand, have not been updated since 2004. These do not specify any environmental considerations, dealing exclusively with the most minimum requirements and classifications used to grade the quality of fruit.

There is thus an opportunity to address this nexus between biodiversity and agriculture through a draft issuance from the DENR and DA which provides for the recognition and certification of biodiversity-friendly agricultural practices in terrestrial and aquatic farms. The joint Administrative Order that seeks to mainstream these practices, particularly around PAs, has yet to be approved (DENR-BMB, 2018), so it remains to be seen whether this might apply to existing plantations, and if any penalties or disincentives will be applied in case of non-compliance.

Thirdly, the review of policies reveals that there is a consistent need to expand the coverage and regulation for PAs. Philippine PAs are not meeting the biodiversity coverage of the Aichi targets (Mallari et al., 2016), which means that habitats and species remain at large risk to being lost to human activity, including climate change effects, agricultural expansion, and its intensification. However, this may be a challenging demand on government resources, and recognizing the acute violence and threats to under-supported forest rangers and protectors, including indigenous peoples, as well as the increasing pressures from agricultural and urban LULUC. While legislation and policy may be tools to afford some protection to species and habitats in the Philippines, it is clear that the interactions between human society’s needs and nature are very complex.

## 5. Conclusions

Agricultural production and the trade of its products are important economic activities at the local and national level, particularly for high-value crops which can generate large revenues. However, the continued increases in agricultural intensification and expansion to grow input-heavy crops such as banana and pineapple have large impacts on biodiversity, particularly for threatened species that are already being affected by habitat loss and the direct impacts of agricultural LULUC.

The Philippines presents a case wherein a domestic policy environment which does not have stringent definitions nor protections for biodiversity in areas where there are interactions between species habitats, PAs, IBAs; furthermore agricultural land is coupled with a political and economic context that emphasizes large-scale and export-oriented trade in high value crops. Because of this, important opportunities to protect dwindling natural habitats and ecosystems are often overlooked.

It is strongly recommended that the global, big-picture view of the nexus of agriculture, trade, and biodiversity is considered in environmental protection laws and programs. This may be done through supply-side interventions, such as regulations in permitting and certification, or by raising awareness to influence market demands.

## Acknowledgments and disclosure statement

AMDO and JNVT declare no conflict of interest. AMDO and JNVT jointly wrote and conducted the analysis for the article. AMDO is funded by UK NERC at their home institution.

## Supplementary information

The Appendix for this article is available at: https://tinyurl.com/y2w68unx.

