## Supplementary Information for "Assessing the impacts of agriculture and its trade on Philippine biodiversity"

**Appendix**

*Table A1. Banana plantations.*

| | | |
|---|---|---|
| **Montemar Foods and AgriVenture Corp.**<br><br>Certificate of Registration of Growers/Farmers issued by the BPI in May 2019 at http://bpi.da.gov.ph/bpi/index.php/sample-levels/national-plant-quarantine-services-division/certificate-of-accreditation-of-farmers-growers/5876-certificate-of-registration-of-growers-farmers-of-montemar-foods-agriventure-corp | | |
| **Surigao del Sur** | Tago Municipality | Barangay Victoria (1) |
| **Vizcaya Plantation Inc.**<br><br>Certificate of Registration of Growers/Farmers issued by the BPI in December 2018 at http://bpi.da.gov.ph/bpi/index.php/sample-levels/national-plant-quarantine-services-division/certificate-of-accreditation-of-farmers-growers/5051-certificate-of-registration-as-growers-farmers-of-vizcaya-plantation-inc | | |
| **Compostela Valley** | Maco Municipality | Barangay Dumlan (1) |
| **Highland Agri-Ventures Inc.**<br><br>Certificate of Registration of Growers/Farmers issued by the BPI in February 2018 at http://bpi.da.gov.ph/bpi/index.php/sample-levels/national-plant-quarantine-services-division/certificate-of-accreditation-of-farmers-growers/5245-certificate-of-accreditation-of-growers-farmers-of-highland-agri-ventures-inc | | |
| **Davao del Sur** | Digos City | Barangay Kapatagan (Purok Mt. Apo, 1) |
| **Lapanday Foods Corp**<br><br>Certificate of Registration of Growers/Farmers issued by the BPI in July 2019, July 2018, October 2018 and November 2018 at http://bpi.da.gov.ph/bpi/index.php/sample-levels/national-plant-quarantine-services-division/certificate-of-accreditation-of-farmers-growers/6250-lapanday-foods-corporation and http://bpi.da.gov.ph/bpi/index.php/sample-levels/national-plant-quarantine-services-division/certificate-of-accreditation-of-farmers-growers/4708-certificate-of-registration-as-growers-farmers-of-lapanday-foods-corporation http://bpi.da.gov.ph/bpi/index.php/sample-levels/national-plant-quarantine-services-division/certificate-of-accreditation-of-farmers-growers/4872-certificate-of-registration-as-growers-farmers-of-lapanday-foods-corporation-3 and http://bpi.da.gov.ph/bpi/index.php/sample-levels/national-plant-quarantine-services-division/certificate-of-accreditation-of-farmers-growers/4308-certificate-of-accreditation-of-growers-farmers-lapanday-foods-corporation-2 | | |
| **Davao del Norte** | Panabo City | Barangay Datu Abdul Dadia (1) |
| | Tagum City | Barangay Madaum (1) |
| **Davao del Sur** | Hagonoy Municipality | Barangay Leling (1)<br><br>Barangay Tulogan (2)<br><br>Barangay Paligue (2)<br>Barangay Crossing (1)<br><br>Barangay Guihing (2) |
| | Padada Municipality | Barangay Sergio Osmena (1) |
| **Davao City** | Buhangin District | Barangay Callawa (1) |
| **Sultan Kudarat** | Lambayong Municipality | Barangay Sadsalan (1) |
| **South Cotabato** | Tampakan Municipality | Unspecified (1) |
| **Agusan del Sur** | Trento Municipality | Barangay Poblacion (1) |

| | | |
|---|---|---|
| **Alfaras Banana Cavendish**<br><br>Certificate of Registration of Growers/Farmers issued by the BPI in July 2019 at http://bpi.da.gov.ph/bpi/index.php/sample-levels/national-plant-quarantine-services-division/certificate-of-accreditation-of-farmers-growers/6242-alfaras-banana-cavendish | | |
| **Davao del Norte** | Tagum City | Barangay Pagsabangan (Purok Bayanihan, 1) |
| **Verde Horizon Agri Ventures Corp.**<br><br>Certificate of Registration of Growers/Farmers issued by the BPI in July 2019 at http://bpi.da.gov.ph/bpi/index.php/sample-levels/national-plant-quarantine-services-division/certificate-of-accreditation-of-farmers-growers/6247-verde-horizon-agri-ventures-corporation | | |
| **Davao del Norte** | Carmen Municipality | Barangay Magsaysay (Purok 4A and Purok 4B, 2) |
| | Tagum City | Barangay Mankilam (Purok Lomboy, 1) |
| **Compostela Valley** | Nabunturan Municipality | Barangay Mainit (Purok 2, 1) |
| | Monkayo Municipality | Barangay Babag (Purok 1, 1) |
| | Mawab Municipality | Barangay Sawangan (Sitio Pagud, 1) |
| **Davao del Sur** | Malalag Municipality | Barangay San Isidro (Purok 6, 1) |
| **Sunseeds Trading Corp.**<br><br>Certificate of Registration of Growers/Farmers issued by the BPI in July 2019 at http://bpi.da.gov.ph/bpi/index.php/sample-levels/national-plant-quarantine-services-division/certificate-of-accreditation-of-farmers-growers/6238-sunseeds-trading-corp | | |
| **Davao del Norte** | Tagum City | Barangay Mankilam (Sitio Magtaya, 1) |
| | Panabo City | Barangay Datu Abdul Dadia (Purok Nangka, 1) |
| **Tagum Agricultural Development Corp. Inc.**<br><br>Certificate of Registration of Growers/Farmers issued by the BPI in March 2019 at http://bpi.da.gov.ph/bpi/index.php/sample-levels/national-plant-quarantine-services-division/certificate-of-accreditation-of-farmers-growers/5524-certificate-of-accreditation-of-growers-farmers-of-tagum-agricultural-development-corporation-inc | | |
| **Davao del Norte** | Panabo City | Barangay A.O. Floirendo (12)<br><br>Barangay Sindaton (Purok 6, 1) |
| | Sto. Tomas Municipality | Barangay Balagunan (1) |
| | Braulio E. Dujali Municipality | Barangay Tanglaw (1) |
| **Davao City** | Toril District | Barangay Marapangi (1) |
| **Ating Lupa Bagong Simula Corp.**<br><br>Certificate of Registration of Growers/Farmers issued by the BPI in December 2018 at http://bpi.da.gov.ph/bpi/index.php/sample-levels/national-plant-quarantine-services-division/certificate-of-accreditation-of-farmers-growers/5044-certificate-of-registration-as-growers-farmers-of-ating-lupa-bagong-simula-albasi-corporation | | |
| **Davao del Norte** | Tagum City | Barangay Mankilam (Purok Maligaya, 1) |
| **Henry Villarosa** | | |

<table>
<tr><td colspan="3">Certificate of Registration of Growers/Farmers issued by the BPI in December 2018 at http://bpi.da.gov.ph/bpi/index.php/sample-levels/national-plant-quarantine-services-division/certificate-of-accreditation-of-farmers-growers/5043-certificate-of-registration-as-growers-farmers-of-henry-villarosa</td></tr>
<tr><td>Davao del Norte</td><td>Tagum City</td><td>Barangay Cuambogan (Purok Mercury, 1)</td></tr>
<tr><td colspan="3">Helon International Trading Corp.<br><br>Certificate of Registration of Growers/Farmers issued by the BPI in October 2018 at http://bpi.da.gov.ph/bpi/index.php/sample-levels/national-plant-quarantine-services-division/certificate-of-accreditation-of-farmers-growers/4693-certificate-of-accreditation-of-growers-farmers-of-helon-international-trading-corp</td></tr>
<tr><td>Davao del Norte</td><td>Tagum City</td><td>Barangay La Filipina (Purok 1, 1)<br><br>Barangay Pagsabangan (1)</td></tr>
<tr><td></td><td>Sto. Tomas Municipality</td><td>Barangay Balagunan (1)<br><br>Barangay Kinamayan (1)<br><br>Barangay Bobongon (2)</td></tr>
<tr><td></td><td>Panabo City</td><td>Barangay Consolacion (2)</td></tr>
<tr><td colspan="3">Domingo Bentulan, Wilfredo Saldua, Marites Cavite, Pelagua Saldua, Jocelyn Villanueva, Dionisio Suraibcay, Shariff Bangoy, Roselier Bangoy, Luisito Mantepio, Allan Ong, Rolando Diaz and Ricky Hao<br><br>Certificate of Registration of Growers/Farmers issued by the BPI in March 2018 http://bpi.da.gov.ph/bpi/index.php/sample-levels/national-plant-quarantine-services-division/certificate-of-accreditation-of-farmers-growers/3934-certificate-of-accreditation-of-growers-farmers-helon-farms-2018</td></tr>
<tr><td>Davao del Norte</td><td>Panabo City</td><td>Barangay Consolacion (2)<br><br>Barangay Upper Licanan (Purok 3, 1)<br><br>Barangay Cangangohan (Purok Lanzones, 1)</td></tr>
<tr><td></td><td>Sto. Tomas Municipality</td><td>Barangay Kinamayan (3, 1 in Purok 2)<br><br>Barangay Bobongon (2)<br><br>Barangay Balagunan (1)</td></tr>
<tr><td></td><td>Tagum City</td><td>Barangay La Filipina (Purok 1, 1)<br><br>Barangay Pagsabangan (1)</td></tr>
<tr><td colspan="3">Al-Mujahidun Agro Resources and Development Inc.<br><br>Certificate of Registration of Growers/Farmers issued by the BPI in June 2019 http://bpi.da.gov.ph/bpi/index.php/sample-levels/national-plant-quarantine-services-division/certificate-of-accreditation-of-farmers-growers/5973-ertificate-of-registration-of-growers-farmers-of-al-mujahidun-agro-resources-development-inc</td></tr>
<tr><td>Maguindanao</td><td>Ampatuan Municipality</td><td>Barangay Salman (1)</td></tr>
<tr><td colspan="3">Delinanas Development Corp.<br><br>Certificate of Registration of Growers/Farmers issued by the BPI in June 2019 and June 2018 at http://bpi.da.gov.ph/bpi/index.php/sample-levels/national-plant-quarantine-services-division/certificate-of-accreditation-of-farmers-growers/5722-certificate-of-registration-of-growers-farmers-of-delinanas-development-corporation and http://bpi.da.gov.ph/bpi/index.php/sample-levels/national-plant-quarantine-services-division/certificate-of-accreditation-of-farmers-growers/4141-certificate-of-accreditation-of-growers-farmers-delinanas-farm-cot-2018</td></tr>
</table>

<table>
<tr><td>Maguindanao</td><td>Datu Abdullah Sangki Municipality</td><td>Barangay Tukanalugong (1)<br><br>Barangay Maranding (1)</td></tr>
<tr><td>Sultan Kudarat</td><td>Lambayong Municipality</td><td>Barangay Seneben (1)<br><br>Barangay Sigayan (1)<br><br>Barangay New Cebu (1)<br><br>Barangay Pimbalayan (1)</td></tr>
<tr><td></td><td>Lutayan Municipality</td><td>Barangay Maindang (2)<br><br>Barangay Mamali (1)</td></tr>
<tr><td>North Cotabato</td><td>Tulunan Municipality</td><td>Barangay Dungos (1)</td></tr>
<tr><td colspan="3">Alip River Development and Export Corporation<br><br>Certificate of Registration of Growers/Farmers issued by the BPI in October 2018 at http://bpi.da.gov.ph/bpi/index.php/sample-levels/national-plant-quarantine-services-division/certificate-of-accreditation-of-farmers-growers/4712-certificate-of-registration-as-growers-farmers-of-alip-river-development-export-corporation-ardexcor</td></tr>
<tr><td>Maguindanao</td><td>Datu Paglas Municipality</td><td>Barangay Alip (1)</td></tr>
<tr><td colspan="3">Al-Sahar Agri-Ventures Inc.<br><br>Certificate of Registration of Growers/Farmers issued by the BPI in October 2018 and January 2018 at http://bpi.da.gov.ph/bpi/index.php/sample-levels/national-plant-quarantine-services-division/certificate-of-accreditation-of-farmers-growers/4704-certificate-of-registration-as-growers-farmers-of-al-sahar-agri-ventures-inc and http://bpi.da.gov.ph/bpi/index.php/sample-levels/national-plant-quarantine-services-division/certificate-of-accreditation-of-farmers-growers/3728-certificate-of-accreditation-of-growers-farmers-al-sahar</td></tr>
<tr><td>Maguindanao</td><td>Talayan Municipality</td><td>Barangay Tamar (1)<br>Barangay Marader (1)</td></tr>
<tr><td colspan="3">La Frutera Inc.<br><br>Certificate of Registration of Growers/Farmers issued by the BPI in October 2018 at http://bpi.da.gov.ph/bpi/index.php/sample-levels/national-plant-quarantine-services-division/certificate-of-accreditation-of-farmers-growers/4679-certificate-of-registration-as-growers-farmers-of-la-frutera-inc</td></tr>
<tr><td>Maguindanao</td><td>Buluan Municipality</td><td>Barangay Siling (1, but does not specify whether this Barangay Upper Siling or Lower Siling)<br><br>Also listed are Talingco (1), Liong (1), Linek (1) and Sawa (1) but these aren’t on the list of Buluan barangays.</td></tr>
<tr><td colspan="3">Eka Salam Agriventures Corp.<br><br>Certificate of Registration of Growers/Farmers issued by the BPI in July 2018 at http://bpi.da.gov.ph/bpi/index.php/sample-levels/national-plant-quarantine-services-division/certificate-of-accreditation-of-farmers-growers/4372-certificate-of-accreditation-of-growers-farmers-eka-salam-farm</td></tr>
<tr><td>Maguindanao</td><td>Ampatuan Municipality</td><td>Barangay Kauran (Sitio Sabaduan, 1)</td></tr>
<tr><td colspan="3">Dole Philippines Inc.<br><br>Certificate of Registration of Growers/Farmers issued by the BPI in May 2019, March 2019 and December 2018 at http://bpi.da.gov.ph/bpi/index.php/sample-levels/national-plant-quarantine-services-division/certificate-of-accreditation-of-farmers-growers/5881-certificate-of-registration-of-growers-farmers-of-dole-philippines-inc</td></tr>
</table>

| | | |
|---|---|---|
| http://bpi.da.gov.ph/bpi/index.php/sample-levels/national-plant-quarantine-services-division/certificate-of-accreditation-of-farmers-growers/5501-certificate-of-accreditation-of-growers-farmers-of-dole-philippines-inc-5, http://bpi.da.gov.ph/bpi/index.php/sample-levels/national-plant-quarantine-services-division/certificate-of-accreditation-of-farmers-growers/5108-certificate-of-accreditation-of-growers-farmers-dole-makilala | | |
| **Sultan Kudarat** | Bagumbayan Municipality | Barangay Kinayao (3) |
| **North Cotabato** | Kidapawan City | Barangay Katipunan (1)<br>Barangay Birada (1)<br>Barangay Perez (1)<br>Barangay Luvimin (1)<br>Barangay Marbel (1)<br>Barangay Meohao (1)<br>Barangay Saguing (1) |
| | Makilala Municipality | Barangay Malabuan (2)<br>Barangay Luna Sur (2, 1 in Sitio Laud)<br>Barangay Garsika (2)<br>Barangay Batasan (Sitio Flortan, 1)<br>Barangay Kisante (1)<br>Barangay Buhay (2)<br>Barangay Buena Vida (1)<br>Barangay Indangan (1) |
| | Magpet Municipality | “Cubanan” is specified, but this is not in the list of barangays |
| **Bukidnon** | Impasug-ong Municipality | Barangay La Fortuna (3)<br>Barangay Impalutao (1)<br>Barangay Cawayan (1)<br>Barangay Capitan Bayong (1)<br>Barangay Kibangan (1)<br>“San Juan” is also specified, but this is not in the list of barangays |
| | Baungon Municipality | Barangay Lingating (2)<br>Barangay Buenavista (1)<br>Barangay Liboran (1) |
| | Sumilao Municipality | Barangay Kisolon (3, all in Sitio Laruk)<br>Barangay Poblacion (2, 1 in Sitio Hanawon) |
| | Talakag Municipality | Barangay San Isidro (2) |

| | | |
|---|---|---|
| | | Barangay Lingi-on (1)<br><br>Barangay Sto. Nino (2)<br><br>Barangay San Antonio (1)<br><br>Barangay Dagumbaan (1) |
| | Lantapan Municipality | Barangay Bantuanon (3)<br><br>Barangay Bugcaon (2, both Sitio Sobsob)<br><br>Barangay Alanib (1)<br><br>Barangay Poblacion (3, 1 in Sitio Babahagon)<br><br>Barangay Cawayan (2) |
| | Malaybalay City | Barangay Casisang (6, 1 in Sitio Sta. Ana, 2 in Sitio Sta. Cruz, 1 in Sitio Regla, 2 in Sitio Gabunan)<br><br>“New Ilocos” is also specified, but this is not in the list of barangays |
| **Misamis Oriental** | Tagoloan Municipality | Barangay Sta. Ana (2) |
| | Claveria Municipality | Barangay Mat-I (1) |
| **5J Agduma Banana Farm**<br><br>Certificate of Registration of Growers/Farmers issued by the BPI in August 2018 at http://bpi.da.gov.ph/bpi/index.php/sample-levels/national-plant-quarantine-services-division/certificate-of-accreditation-of-farmers-growers/4450-certificate-of-accreditation-of-growers-farmers-of-5j-agduma-banana-farm | | |
| **Sultan Kudarat** | Lambayong Municpality | Barangay Midtapok (1) |
| **Medalla Farm**<br><br>Certificate of Registration of Growers/Farmers issued by the BPI in April 2018 at http://bpi.da.gov.ph/bpi/index.php/sample-levels/national-plant-quarantine-services-division/certificate-of-accreditation-of-farmers-growers/4032-certificate-of-accreditation-of-growers-farmers-medalla | | |
| **Sultan Kudarat** | Bagumbayan Municipality | Barangay Biwang (Sitio Crismor, 1) |
| **Mindanao Agritraders Inc.**<br><br>Certificate of Registration of Growers/Farmers issued by the BPI in July 2018 and October 2018 at http://bpi.da.gov.ph/bpi/index.php/sample-levels/national-plant-quarantine-services-division/certificate-of-accreditation-of-farmers-growers/4691-certificate-of-accreditation-of-growers-farmers-of-mindanao-agritraders-inc-2 http://bpi.da.gov.ph/bpi/index.php/sample-levels/national-plant-quarantine-services-division/certificate-of-accreditation-of-farmers-growers/4692-certificate-of-accreditation-of-growers-farmers-of-mindanao-agritraders-inc-3 http://bpi.da.gov.ph/bpi/index.php/sample-levels/national-plant-quarantine-services-division/certificate-of-accreditation-of-farmers-growers/4305-certificate-of-accreditation-of-growers-farmers-mindanao-agritraders-inc and http://bpi.da.gov.ph/bpi/index.php/sample-levels/national-plant-quarantine-services-division/certificate-of-accreditation-of-package-facility/4304-certificate-of-accreditation-of-packing-facility-mindanao-agritraders-inc | | |
| **Bukidnon** | Malaybalay City | Barangay Cabangahan (1) |
| | Lantapan Municipality | Barangay Bugcaon (1) |
| **Agusan del Norte** | Cabadbaran City | Barangay Soriano (2) |

| | | |
|---|---|---|
| **Manupali Agri Development Corp.**<br><br>Certificate of Registration of Growers/Farmers issued by the BPI in April 2019 at http://bpi.da.gov.ph/bpi/index.php/sample-levels/national-plant-quarantine-services-division/certificate-of-accreditation-of-farmers-growers/5622-certificate-of-accreditation-of-growers-farmers-of-manupali-agri-development-corporation | | |
| **Bukidnon** | Valencia City | Certificate specifies “Dabongdabong,” but this is not among the listed barangays |
| **Sumifru Agricultural Development Inc.**<br><br>Certificate of Registration of Growers/Farmers issued by the BPI in May 2019 at http://bpi.da.gov.ph/bpi/index.php/sample-levels/national-plant-quarantine-services-division/certificate-of-accreditation-of-farmers-growers/5855-certificate-of-registration-of-growers-farmers-of-sumifru-agricultural-development-inc | | |
| **Bukidnon** | Valencia City | Barangay Guinoyuran (1)<br><br>Barangay Lurugan (1) |
| **Grand Terrain Corporation**<br><br>Certificate of Registration of Growers/Farmers issued by the BPI in October 2018 at http://bpi.da.gov.ph/bpi/index.php/sample-levels/national-plant-quarantine-services-division/certificate-of-accreditation-of-farmers-growers/5109-certificate-of-accreditation-of-growers-farmers-grand-terrain-2018 | | |
| **North Cotabato** | Arakan Municipality | Barangay Meocan (1) |
| **SLM Agri Venture Corp.**<br><br>Certificate of Registration of Growers/Farmers issued by the BPI in May 2019 at http://bpi.da.gov.ph/bpi/index.php/sample-levels/national-plant-quarantine-services-division/certificate-of-accreditation-of-farmers-growers/5748-certificate-of-registration-of-growers-farmers-of-slm-agri-venture-corp | | |
| **Agusan del Sur** | Loreto Municipality | Barangay Nueva Gracia (1) |
| **Xinyu Fruits Agricultural Corp**<br><br>Certificate of Registration of Growers/Farmers issued by the BPI in July 2019 at http://bpi.da.gov.ph/bpi/index.php/sample-levels/national-plant-quarantine-services-division/certificate-of-accreditation-of-farmers-growers/6244-xinyu-fruits-agricultural-corporation | | |
| **Davao Oriental** | Lupon Municipality | Barangay Magsaysay (Purok Marang, 1) |
| **ANFLO Banana Corporation**<br><br>Certificate of Registration of Growers/Farmers issued by the BPI in March 2019 at http://bpi.da.gov.ph/bpi/index.php/sample-levels/national-plant-quarantine-services-division/certificate-of-accreditation-of-farmers-growers/5530-certificate-of-accreditation-of-growers-farmers-of-anflo-banana-corporation | | |
| **Davao Oriental** | Lupon Municipality | Barangay Limbahan (1) |
| **JPP Fresh Produce Corp.**<br><br>Certificate of Registration of Growers/Farmers issued by the BPI in December 2018 at http://bpi.da.gov.ph/bpi/index.php/sample-levels/national-plant-quarantine-services-division/certificate-of- | | |

| | | |
|---|---|---|
| accreditation-of-farmers-growers/5059-certificate-of-registration-as-growers-farmers-of-jpp-fresh-produce-corporation | | |
| **Davao Oriental** | Governor Generoso Municipality | Barangay Tiblawan (1) |
| **Dana Fresh Fruits Corp.**<br><br>Certificate of Registration of Growers/Farmers issued by the BPI in December 2018 | | |
| **Davao Oriental** | Banaybanay Municipality | Barangay Calubihan (Purok 9, 1) |
| **Sunnjef Plantation Inc.**<br><br>Certificate of Registration of Growers/Farmers issued by the BPI in January 2019 at http://bpi.da.gov.ph/bpi/index.php/sample-levels/national-plant-quarantine-services-division/certificate-of-accreditation-of-farmers-growers/5137-certificate-of-accreditation-of-growers-farmers-of-sunnjef-plantation-inc | | |
| **North Cotabato** | Makilala Municipality | Barangay Taluntalunan (1)<br><br>Also specifies a “Sta. Cruz, Laguda” but this is not one of the listed barangays |

| | | |
|---|---|---|
| **Sumifru Philippines Corporation**<br><br>(Bureau of Plant Industry Certificate of Registration of Growers/Farmers, issued May 2019 at http://bpi.da.gov.ph/bpi/index.php/sample-levels/national-plant-quarantine-services-division/certificate-of-accreditation-of-farmers-growers/6418-sumifru-philippines-corporation) | | |
| **Davao City** | | |
| | Toril District | Barangay Tungkalan (3)<br><br>Barangay Tibuloy (2) |
| | Tugbok District | Barangay Manuel Guianga (2) |
| | Calinan District | Barangay Tamayong (2)<br><br>Barangay Subasta (1)<br><br>Barangay Dacudao (1) |
| | Baguio District | Barangay Tambobong (2) |
| | Marilog District | Barangay Salaysay (4) |
| **Davao del Norte** | | |
| | Sto. Tomas Municipality | Barangay Casig-ang (1)<br><br>Barangay Kinamayan (1) |

| | | |
|---|---|---|
| | Tagum City | Barangay La Filipina (1) |
| | New Corella Municipality | Barangay San Roque (7) |
| **Compostela Valley** | Compostela Municipality | Barangay Osmeña (2)<br><br>Barangay Mangayon (2)<br><br>Barangay New Alegria (1) |
| | Laak Municipality | Barangay Kapatagan (3, 2 specified in Purok 3)<br><br>Barangay Amor Cruz (2, 1 each in Puroks 1 and 6)<br><br>Barangay Kaligutan (4, 1 each in Puroks 3, 5 and 8, and 1 specifying Barangays Kaligutan and Kilagting) |
| | Mawab Municipality | Barangay Lower Guisok (7, all in Purok 10) |
| **Dole Philippines Inc.**<br><br>(Bureau of Plant Industry Certificate of Registration of Growers/Farmers, issued July 2019 at http://bpi.da.gov.ph/bpi/index.php/sample-levels/national-plant-quarantine-services-division/certificate-of-accreditation-of-farmers-growers/6278-dole-philippines-inc-5, http://bpi.da.gov.ph/bpi/index.php/sample-levels/national-plant-quarantine-services-division/certificate-of-accreditation-of-farmers-growers/6277-dole-philippines-inc-4, http://bpi.da.gov.ph/bpi/index.php/sample-levels/national-plant-quarantine-services-division/certificate-of-accreditation-of-farmers-growers/6269-dole-philippines-inc-3, http://bpi.da.gov.ph/bpi/index.php/sample-levels/national-plant-quarantine-services-division/certificate-of-accreditation-of-farmers-growers/6268-dole-philippines-inc-2, http://bpi.da.gov.ph/bpi/index.php/sample-levels/national-plant-quarantine-services-division/certificate-of-accreditation-of-farmers-growers/6267-dole-philippines-inc) | | |
| **Davao City** | Calinan District | Barangay Tamayong (1)<br><br>Barangay Wangan (1) |
| | Baguio District | Barangay Malagos (1)<br><br>Barangay Gumalang (1)<br><br>Barangay Carmen (1) |
| | Tugbok District | Barangay Talandang (1) |
| **Davao del Norte** | Panabo City | Barangay Kasilak (2, also specifies Buenaventura, could be the owner's name or the Sitio?)<br><br>Barangay Little Panay (2, one specifies Magarang)<br><br>Barangay Datu Abdul Dadia (1, also specifies Ranain) |

| | | |
|---|---|---|
| | | Barangay Dapco (3, also specifies Bananeros and DUSGROW)<br><br>Barangay Katipunan (3, specifies Icasas and Ventic)<br><br>Barangay Manay (1)<br><br>Barangay Upper Licanan (1)<br><br>Others specify “Bandong”, “Survivor” and “Mabuhay” in Panabo City, but these don’t seem to be barangay names. There is a Barangay Mabuhay in Carmen Municipality, but it is unclear if it is the same one. |
| | Asuncion Municipality | Barangay Magatos (1)<br><br>Barangay Camoning (1)<br><br>Barangay Buclad (1)<br><br>One more specifies “Lawang” in Asuncion, but this doesn’t seem to be a barangay name |
| | Kapalong Municipality | Barangay Sampao (2)<br><br>Barangay Pag-asa (4)<br><br>Barangay Tiburcia (1)<br><br>Barangay Luna (1)<br><br>One more specifies “San Jose” in Kapalong, but this doesn’t seem to be a barangay name |
| | Carmen Municipality | Barangay Alejal (4, 1 each issued to DEFARBEMCO, Godifarba and Dagondon). DEFARBEMCO is listed a second time, but no barangay is given.<br><br>Barangay Magsaysay (1)<br><br>Barangay Tubod (1, issued to BSBG)<br><br>Also listed are Almaca and SEARBEMCO, but no barangay is given. One more specifies “Siak” in Carmen, but this doesn’t seem to be a barangay name |
| | Sto. Tomas Municipality | Barangay Lunga-og (2)<br><br>Barangay Kinamayan (1)<br><br>Barangay Talomo (1)<br><br>Barangay Kimamon (1)<br><br>Barangay La Libertad (1)<br><br>Barangay Balagunan (1, issued to Sta. Lucia) |

| | | |
|---|---|---|
| | | Also listed is DFI, but no barangay is given. |
| | New Corella Municipality | Barangay Mesaoy (1)<br><br>Barangay Masagana (1) |
| **Compostela Valley** | Laak Municipality | Barangay Aguinaldo (1, specifies Purok 1) |
| | Maragusan Municipality | Barangay New Albay (1)<br><br>Barangay Pamintaran (1)<br><br>Barangay New Katipunan (2)<br><br>Barangay New Panay (1)<br><br>Barangay Magcagong (1)<br><br>Barangay Mauswagon (1)<br><br>Barangay Tupaz (1)<br><br>Barangay Lahi (1)<br><br>Barangay Mapawa (2)<br><br>Barangay Poblacion (1)<br><br>One more specifies “Kasilak” in Maragusan, but this doesn’t seem to be a barangay name |
| | Pantukan Municipality | Barangay Araibo |
| **Davao del Sur** | Hagonoy Municipality | Barangay Sinayawan (1) |
| | Digos City | Barangay Matti (1)<br><br>Barangay Kapatagan (1) |
| **Panombon Agri-Ventures Inc.**<br><br>(Bureau of Plant Industry Certificate of Registration of Growers/Farmers, issued July 2019 at http://bpi.da.gov.ph/bpi/index.php/sample-levels/national-plant-quarantine-services-division/certificate-of-accreditation-of-farmers-growers/6257-panombon-agri-ventures-inc | | |
| **Davao Oriental** | Mati City | Barangay Don Enrique Lopez |
| **Happy Bananas Agro-Enterprise**<br><br>(Bureau of Plant Industry Certificate of Registration of Growers/Farmers, issued July 2019 at http://bpi.da.gov.ph/bpi/index.php/sample-levels/national-plant-quarantine-services-division/certificate-of-accreditation-of-farmers-growers/6264-happy-bananas-agro-enterprise) | | |

| | | |
|---|---|---|
| **North Cotabato** | Tulunan Municipality | Barangay Dungos (1) |
| **Arnold Miranda Farm**<br><br>(Bureau of Plant Industry Certificate of Registration of Growers/Farmers, issued July 2019 at http://bpi.da.gov.ph/bpi/index.php/sample-levels/national-plant-quarantine-services-division/certificate-of-accreditation-of-farmers-growers/6265-arnold-miranda-farm) | | |
| **North Cotabato** | Tulunan Municipality | Barangay Tambac (1) |
| **Jacinto Tadiaque**<br><br>(Bureau of Plant Industry Certificate of Registration of Growers/Farmers, issued August 2019 at http://bpi.da.gov.ph/bpi/index.php/sample-levels/national-plant-quarantine-services-division/certificate-of-accreditation-of-farmers-growers/6397-jacinto-tadiaque) | | |
| **Davao del Norte** | Santo Tomas Municipality | Barangay New Visayas (Purok Narra, 1) |
| **Ramonita Tadiaque**<br><br>(Bureau of Plant Industry Certificate of Registration of Growers/Farmers, issued August 2019 at http://bpi.da.gov.ph/bpi/index.php/sample-levels/national-plant-quarantine-services-division/certificate-of-accreditation-of-farmers-growers/6381-ramonita-tadiaque) | | |
| **Davao del Norte** | Santo Tomas Municipality | Barangay New Visayas (1) |
| **Ronald Allan Matute Farm**<br><br>(Bureau of Plant Industry Certificate of Registration of Growers/Farmers, issued September 2019 at<br><br>http://bpi.da.gov.ph/bpi/index.php/sample-levels/national-plant-quarantine-services-division/certificate-of-accreditation-of-farmers-growers/6624-ronald-allan-matute-farm) | | |
| **Compostela Valley** | Not specified | Not specified (1) |
| **Granada Banana Farm**<br><br>(Bureau of Plant Industry Certificate of Registration of Growers/Farmers, issued July 2019 at http://bpi.da.gov.ph/bpi/index.php/sample-levels/national-plant-quarantine-services-division/certificate-of-accreditation-of-farmers-growers/6496-granada-banana-farm | | |
| **North Cotabato** | Tulunan Municipality | Barangay La Esperanza (1) |
| **Villamor Banana Farm**<br><br>(Bureau of Plant Industry Certificate of Registration of Growers/Farmers, issued August 2019 at http://bpi.da.gov.ph/bpi/index.php/sample-levels/national-plant-quarantine-services-division/certificate-of-accreditation-of-farmers-growers/6494-villamor-banana-farm | | |

| | | |
|---|---|---|
| **North Cotabato** | Tulunan Municipality | Barangay Damawato (1) |
| **Lemana Banana Farm**<br><br>(Bureau of Plant Industry Certificate of Registration of Growers/Farmers, issued August 2019 at http://bpi.da.gov.ph/bpi/index.php/sample-levels/national-plant-quarantine-services-division/certificate-of-accreditation-of-farmers-growers/6492-lemana-banana-farm) | | |
| **North Cotabato** | Tulunan Municipality | Barangay Bual (Sitio Casalan, 1) |
| **Sorilla (Farm 80) Banana Farm**<br><br>(Bureau of Plant Industry Certificate of Registration of Growers/Farmers, issued August 2019 at http://bpi.da.gov.ph/bpi/index.php/sample-levels/national-plant-quarantine-services-division/certificate-of-accreditation-of-farmers-growers/6491-sorilla-farm-80-banana-farm) | | |
| **North Cotabato** | Tulunan Municipality | Barangay Sibsib (1) |
| **EOS Mindatrade International Corporation**<br><br>(Bureau of Plant Industry Certificate of Registration of Growers/Farmers, issued August 2019 at http://bpi.da.gov.ph/bpi/index.php/sample-levels/national-plant-quarantine-services-division/certificate-of-accreditation-of-farmers-growers/6400-eos-mindatrade-international-corporation) | | |
| **Davao City** | Tugbok District | Barangay Manuel Guianga (1) |
| **Compostela Valley** | New Bataan Municipality (website says “New Bantacan,” this might be closest?) | Not specified (1) |
| **Davao del Norte** | Kapalong Municipality | Barangay Pag-asa (4, 1 in Purok 4)<br><br>Barangay Maniki (2)<br><br>Barangay Luna (1)<br><br>Balmar is specified, but no barangay is given |
| | Carmen Municipality | Barangay Sto. Niño (Purok 7, 1)<br><br>Barangay Cebulano (2)<br><br>Barangay Tubod (3, 1 in Purok 3B) |
| | Asuncion Municipality | Barangay Napungas (1)<br><br>Barangay New Bantayan (1) |
| | Santo Tomas Municipality | Barangay Lunga-og (3) |

| | | |
|---|---|---|
| | | Barangay La Libertad (2, 1 in Purok 7)<br>Barangay Casig-ang (1)<br>Barangay Talomo (2)<br>Barangay Balagunan (1) |
| **Avante Agri Products Inc.**<br>(Bureau of Plant Industry Certificate of Registration of Growers/Farmers, issued September 2019 at http://bpi.da.gov.ph/bpi/index.php/sample-levels/national-plant-quarantine-services-division/certificate-of-accreditation-of-farmers-growers/6647-avante-agri-products-philippines-inc) | | |
| **Davao del Norte** | Tagum City | Barangay Pagsabangan (Barrio Mankilam, 2) |
| **Impreza Fruit Trading**<br>(Bureau of Plant Industry Certificate of Registration of Growers/Farmers, issued September 2019 at http://bpi.da.gov.ph/bpi/index.php/sample-levels/national-plant-quarantine-services-division/certificate-of-accreditation-of-farmers-growers/6645-impreza-fruit-trading) | | |
| **Davao del Norte** | Sto. Tomas Municipality | Barangay Bobongon (2, 1 in Purok 2) |
| **Compostela Valley** | Compostela Municipality | Barangay Tamia (1) |
| **823 Packing Plant**<br>(Bureau of Plant Industry Certificate of Registration of Growers/Farmers, issued September 2019 at http://bpi.da.gov.ph/bpi/index.php/sample-levels/national-plant-quarantine-services-division/certificate-of-accreditation-of-farmers-growers/6644-823-packing-plant) | | |
| **Davao del Norte** | Sto. Tomas Municipality | Barangay Balagunan (Purok 1, 1)<br>Barangay Kinamayan (1)<br>Barangay Lungaog (Purok Rizal, 1)<br>Barangay Tibal-og (Purok 5, 1) |
| **Nader and Ebrahim S/O Hassan Philippines Inc.**<br>(Bureau of Plant Industry Certificate of Registration of Growers/Farmers, issued September 2019 at http://bpi.da.gov.ph/bpi/index.php/sample-levels/national-plant-quarantine-services-division/certificate-of-accreditation-of-farmers-growers/6642-nader-ebrahim-s-o-hassan-philippines-inc) | | |
| **Davao del Norte** | Sto. Tomas Municipality | Barangay Kinamayan (Purok 6, 1) |
| | Tagum City | Barangay Cuambogan (Purok Mercury, 1) |

| | | |
|---|---|---|
| | New Corella Municipality | Barangay Mesaoy (Purok 14, 1) |
| | Carmen Municipality | Barangay San Isidro (1) |
| **Compostela Valley** | Maco Municipality | Barangay Hijo (1) |
| **Tortuga Valley Plantation Inc.**<br><br>**(Bureau of Plant Industry Certificate of Registration of Growers/Farmers, issued September 2019 at**<br><br>**http://bpi.da.gov.ph/bpi/index.php/sample-levels/national-plant-quarantine-services-division/certificate-of-accreditation-of-farmers-growers/6640-tortuga-valley-plantation-inc** | | |
| **Davao del Sur** | Digos City | Barangay Igpit (1) |
| **Kapatagan Banana Growers Cooperative**<br><br>**(Bureau of Plant Industry Certificate of Registration of Growers/Farmers, issued September 2019 at http://bpi.da.gov.ph/bpi/index.php/sample-levels/national-plant-quarantine-services-division/certificate-of-accreditation-of-farmers-growers/6637-kapatagan-banana-growers-coop** | | |
| **Davao del Sur** | Digos City | Barangay Kapatagan (Purok Mauswagon, 1) |

*Table A2. Pineapple plantations.*

| | | |
|---|---|---|
| **Sumifru Philippines Corp.**<br><br>Certificate of Registration of Growers/Farmers issued by the BPI in January 2019 and May 2019 at http://bpi.da.gov.ph/bpi/index.php/sample-levels/national-plant-quarantine-services-division/certificate-of-accreditation-of-farmers-growers/5082-certificate-of-accreditation-of-growers-farmers-of-sumifru-philippines-corporation-2 and http://bpi.da.gov.ph/bpi/index.php/sample-levels/national-plant-quarantine-services-division/certificate-of-accreditation-of-farmers-growers/5857-certificate-of-registration-of-growers-farmers-of-sumifru-philippines-corporation-6 | | |
| **Bukidnon** | Valencia City | Barangay Barobo (1)<br><br>Barangay Lourdes (1)<br><br>Barangay San Carlos (1)<br><br>Barangay Lurugan (1) |
| | Malaybalay City | Barangay Magsaysay (2)<br><br>Barangay Cabangahan (1) |
| | Lantapan Municipality | Barangay Bugcaon (1)<br><br>Barangay Bantuanon (1)<br><br>Barangay Kulasihan (1)<br><br>Unspecified Barangay (1) |

| | | |
|---|---|---|
| | Cabanglasan Municipality | Barangay Iba (1) |
| **S&N Fruits Corporation**<br><br>Certificate of Registration of Growers/Farmers issued by the BPI in January 2019 at http://bpi.da.gov.ph/bpi/index.php/sample-levels/national-plant-quarantine-services-division/certificate-of-accreditation-of-farmers-growers/5081-certificate-of-accreditation-of-growers-farmers-of-s-n-fruits-corporation | | |
| **South Cotabato** | Tampakan Municipality | Barangay Lambayong (1) |

| | | |
|---|---|---|
| **Dole Philippines Inc.**<br><br>**(Bureau of Plant Industry Certificate of Registration of Growers/Farmers, issued August 2019 at http://bpi.da.gov.ph/bpi/index.php/sample-levels/national-plant-quarantine-services-division/certificate-of-accreditation-of-farmers-growers/6489-dole-philippines-inc-8)** | | |
| **Sultan Kudarat** | Columbio Municipality | Barangay Polomolok (1) |
| **Wao Development Corporation**<br><br>**(Bureau of Plant Industry Certificate of Registration of Growers/Farmers, issued July 2019 at http://bpi.da.gov.ph/bpi/index.php/sample-levels/national-plant-quarantine-services-division/certificate-of-accreditation-of-farmers-growers/6110-certificate-of-registration-of-growers-farmers-of-wao-development-corporation** | | |
| **Lanao del Sur** | Wao Municipality | Barangay Manila Group (1) |
| **JPY Farms and Agri Supply**<br><br>**(Bureau of Plant Industry Certificate of Registration of Growers/Farmers, issued May 2019 at http://bpi.da.gov.ph/bpi/index.php/sample-levels/national-plant-quarantine-services-division/certificate-of-accreditation-of-farmers-growers/5737-certificate-of-registration-of-growers-farmers-of-jpy-farms-and-agri-supply** | | |
| **Lanao del Sur** | Wao Municipality | Barangay Kabatangan (1) |
| **Southern Philippine Fresh Fruits Corporation**<br><br>**(Bureau of Plant Industry Certificate of Registration of Growers/Farmers, issued April 2019 at http://bpi.da.gov.ph/bpi/index.php/sample-levels/national-plant-quarantine-services-division/certificate-of-accreditation-of-farmers-growers/5630-certificate-of-accreditation-of-growers-farmers-of-southern-philippine-fresh-fruits-corp** | | |
| **South Cotabato** | Polomolok Municipality | Barangay Sulit (1) |
| **Nursery Farms**<br><br>**(Bureau of Plant Industry Certificate of Registration of Growers/Farmers, issued February 2019 at http://bpi.da.gov.ph/bpi/index.php/sample-levels/national-plant-quarantine-services-division/certificate-of-accreditation-of-farmers-growers/5165-certificate-of-accreditation-of-growers-farmers-of-nursery-farms** | | |
| **South Cotabato** | Polomolok Municipality | Barangay Polo (1) |

| | | |
|---|---|---|
| **Davao Agricultural Ventures Corporation**<br><br>**(Bureau of Plant Industry Certificate of Registration of Growers/Farmers, issued January 2019 and May 2019 at http://bpi.da.gov.ph/bpi/index.php/sample-levels/national-plant-quarantine-services-division/certificate-of-accreditation-of-farmers-growers/5140-certificate-of-accreditation-of-growers-farmers-of-davao-agricultural-ventures-corporation-2 and http://bpi.da.gov.ph/bpi/index.php/sample-levels/national-plant-quarantine-services-division/certificate-of-accreditation-of-farmers-growers/5848-certificate-of-registration-of-growers-farmers-of-davao-agricultural-ventures-corp)** | | |
| **Davao City** | Calinan District | Barangay Cawayan (1) |
| **Bukidnon** | Don Carlos Municipality | Barangay San Nicolas (1) |
| | Quezon Municipality | Barangay Merangerang (1) |
| **Natures Fresh Pineapple Inc.**<br><br>**(Bureau of Plant Industry Certificate of Registration of Growers/Farmers, issued May 2019 at http://bpi.da.gov.ph/bpi/index.php/sample-levels/national-plant-quarantine-services-division/certificate-of-accreditation-of-farmers-growers/5708-certificate-of-accreditation-of-growers-farmers-of-nature-s-fresh-pineapple-inc)** | | |
| **Bukidnon** | Malaybalay City | Barangay Sto. Niño and Barangay San Jose (1)<br><br>Barangay Magsaysay (1)<br><br>Barangay Aglayan (1)<br><br>Sitio Patag in Barangay Bugcaon is listed twice, under Malaybalay City and Lantapan Municipality. There is no Barangay Bugcaon in Malaybalay City, so it might be mislabelled. |
| | Maramag Municipality | Barangay Panalsalan (1) |
| | Lantapan Municipality | Barangay Bugcaon (Sitio Patag, 1) |
| **Mt. Kitanglad Agri Development Corporation**<br><br>**(Bureau of Plant Industry Certificate of Registration of Growers/Farmers, issued April 2019 at http://bpi.da.gov.ph/bpi/index.php/sample-levels/national-plant-quarantine-services-division/certificate-of-accreditation-of-farmers-growers/5638-certificate-of-accreditation-of-growers-farmers-of-mt-kitanglad-agri-development-corporation)** | | |
| **Bukidnon** | Valencia City | Barangay Lurugan (1) |
| **South Bukidnon Fresh Trading**<br><br>**(Bureau of Plant Industry Certificate of Registration of Growers/Farmers, issued April 2019 at http://bpi.da.gov.ph/bpi/index.php/sample-levels/national-plant-quarantine-services-division/certificate-of-accreditation-of-farmers-growers/5636-certificate-of-accreditation-of-growers-farmers-of-south-bukidnon-fresh-trading-inc-3)** | | |

| | | |
|---|---|---|
| **Bukidnon** | Don Carlos Municipality | Barangay not specified (1) |
| | Libona Municipality | Barangays not specified (2, except to say West Area and North Area) |
| | Manolo Fortich Municipality | Barangay not specified (1) |
| | Sumilao Municipality | Barangay not specified (1) |
| | Impasug-ong Municipality | Barangay not specified (1) |
| | Malaybalay City | Barangay not specified (1) |
| | Lantapan Municipality | Barangay not specified (1) |
| | Quezon Municipality | Barangay not specified (1) |
| | Maramag Municipality | Barangay not specified (1) |
| | Pangantucan Municipality | Barangay not specified (1) |
| **PhilPack**<br><br>**(Bureau of Plant Industry Certificate of Registration of Growers/Farmers, issued April 2019 at [http://bpi.da.gov.ph/bpi/index.php/sample-levels/national-plant-quarantine-services-division/certificate-of-accreditation-of-farmers-growers/5632-certificate-of-accreditation-of-growers-farmers-of-philpack-2](http://bpi.da.gov.ph/bpi/index.php/sample-levels/national-plant-quarantine-services-division/certificate-of-accreditation-of-farmers-growers/5632-certificate-of-accreditation-of-growers-farmers-of-philpack-2)** | | |
| **Bukidnon** | Don Carlos Municipality | Barangay not specified (1) |
| | Libona Municipality | Barangays not specified (2) |
| | Manolo Fortich Municipality | Barangay not specified (1) |
| | Sumilao Municipality | Barangay not specified (1) |
| | Quezon Municipality | Barangay not specified (1) |
| | Malaybalay City | Barangay not specified (1) |

| | | |
|---|---|---|
| | Impasug-ong Municipality | Barangay not specified (1) |
| | Maramag Municipality | Barangay not specified (1) |
| | Lantapan Municipality | Barangay not specified (1) |
| | Pangantucan Municipality | Barangay not specified (1) |

*Table A3. Potential overlaps with buffer zones and Protected Areas.*

| PA | Possible Overlaps (Area) | Possible Overlaps (Plantation) |
|---|---|---|
| **Mt. Balatukan Range Natural Park**<br><br>Proclamation 1249 s. 2007 and E-NIPAS Coordinates at: https://www.officialgazette.gov.ph/2007/03/06/proclamation-no-1249-s-2007/ | **Misamis Oriental**<br><br>• Municipality of Claveria | Dole Philippines Inc. (banana) |
| **Mt. Hamiguitan Range Wildlife Sanctuary**<br><br>RA 9303 (2004)<br>Coordinates at: https://www.lawphil.net/statutes/repacts/ra2004/ra_9303_2004.html | **Davao Oriental**<br><br>• Municipality of Governor Generoso<br>• Mati City | JPP Fresh Produce Corp. - Gov. Generoso (banana)<br><br>Panombon Agri-ventures Inc. - Mati (banana) |
| **Mt. Kitanglad Range Protected Area**<br><br>RA 8978 (2000)<br>Coordinates at: https://thecorpusjuris.com/legislative/republic-acts/ra-no-8978.php | **Bukidnon**<br><br>• Malaybalay City<br>• Talakag Municipality<br>• Lantapan Municipality<br>• Impasug-ong Municipality<br>• Libona Municipality<br>• Manolo Fortich Municipality | Dole Philippines Inc. - Talakag, Malaybalay, Lantapan, Impasug-ong (banana)<br><br>Mindanao Agritraders Inc. - Malaybalay, Lantapan (banana)<br><br>Sumifru Philippines Corp. - Lantapan, Malaybalay (pineapple)<br><br>South Bukidnon Fresh Trading - Manolo Fortich, Libona, Malaybalay, Impasug-ong, Lantapan, Maramag, Pangantucan (pineapple)<br><br>PhilPack - - Manolo Fortich, Libona, Malaybalay, Impasug-ong, Lantapan, Maramag, Pangantucan (pineapple)<br><br>Nature's Fresh Pineapple - Maramag (pineapple) |
| **Aliwagwag Protected Landscape**<br><br>Proclamation 139 s. 2011 and E-NIPAS | **Compostela Valley**<br><br>• Compostela Municipality | Sumifru Philippines Corp. (banana)<br><br>Impreza Fruit Trading (banana) |

| | | |
|---|---|---|
| Coordinates at<br>https://www.officialgazette.gov.ph/2011/04/05/proclamation-no-139/ | | |
| **Mt. Apo Natural Park**<br><br>RA 9237 (2004)<br>Coordinates at:<br>https://www.officialgazette.gov.ph/2004/02/03/republic-act-no-9237/ | **North Cotabato**<br><br>• Kidapawan City<br>• Makilala Municipality<br>• Magpet Municipality<br><br>**Davao del Sur**<br><br>• Digos City<br><br>**Davao City** | Dole Philippines Inc. - Magpet, Kidapawan, Makilala, Digos, Davao City (banana)<br><br>Sunnjef Plantation Inc. - Makilala (banana)<br><br>Kapatagan Banana Growers Cooperative - Digos (banana)<br><br>Tortuga Valley Plantation - Digos (banana)<br><br>Lapanday Food Corp. - Davao City (banana)<br><br>Tagum Agricultural Development Corp. - Davao City (banana)<br><br>Sumifru Philippines Corp. - Davao City (banana)<br><br>EOS Mindatrade International Corp. - Davao City (banana)<br><br>Davao Agriventures Corp. - Davao City (pineapple)<br><br>According to DENR Region 11, the PA covers Barangays Goma, Binaton, Balabag and Kapatagan in Digos City, and 2 plantations are in Barangay Kapatagan. In Davao City, there might be overlaps in Barangays Tamayong, Manuel Guianga and Tungkalan. In Makilala, there might be overlaps in Barangays Buhay, Batasan and Buena Vida. In Kidapawan, there might be an overlap in Barangay Perez. http://r11.denr.gov.ph/index.php/e-library/manual-of-land-disposition/index.php?option=com_content&view=article&id=339:list-of-protected-areas&catid=100:statistical-data |
| **Mainit Hot Springs National Park**<br><br>Proclamation 320 s. 2000 and E-NIPAS<br>Coordinates at:<br>https://www.officialgazette.gov.ph/2000/05/31/proclamation-no-320-s-2000/ | **Compostela Valley**<br><br>• Nabunturan Municipality | Verde Horizon Agri-ventures Corp. (banana)<br><br>According to DENR Region 11, the PA covers Barangays Mainit and Bukal, and the plantation is also in Barangay Mainit (http://r11.denr.gov.ph/index.php/e-library/manual-of-land-disposition/index.php?option=com_content&view=article&id=339:list-of-protected-areas&catid=100:statistical-data) |

| | | |
|---|---|---|
| **Mati Protected Landscape**<br><br>Proclamation 912 s. 2005 and E-NIPAS Coordinates at: https://www.officialgazette.gov.ph/2005/09/06/proclamation-no-912-s-2005/ | **Davao Oriental**<br><br>• Mati City | Panombon Agri-ventures Inc. (banana)<br><br>According to DENR Region 11, the PA covers Barangays Culian. Sainz and Sudlon, and the plantation is in Barangay Don Enrique Lopez (http://r11.denr.gov.ph/index.php/e-library/manual-of-land-disposition/index.php?option=com_content&view=article&id=339:list-of-protected-areas&catid=100:statistical-data) |
| **Mt. Kalatungan Range Natural Park**<br><br>Proclamation 305 s. 2000 and E-NIPAS Coordinates at: https://www.officialgazette.gov.ph/2000/05/05/proclamation-no-305-s-2000/ | **Bukidnon**<br><br>• Valencia City<br>• Talakag Municipality<br>• Maramag Municipality<br>• Pangantucan Municipality | Dole Philippines Inc. - Talakag, (banana)<br><br>Manupali Agri Development Co. - Valencia (banana)<br><br>Sumifru Agri Development Inc. - Valencia (banana)<br><br>Sumifru Philippines Corp. - Valencia(pineapple)<br><br>South Bukidnon Fresh Trading - Maramag, Pangantucan (pineapple)<br><br>PhilPack - - Manolo Fortich, Libona, Maramag, Pangantucan (pineapple)<br><br>Nature's Fresh Pineapple - Maramag (pineapple) |
